# Dual expression of Atoh1 and Ikzf2 promotes transformation of adult cochlear supporting cells into outer hair cells

**DOI:** 10.1101/2021.01.21.427665

**Authors:** Suhong Sun, Shuting Li, Zhengnan Luo, Minhui Ren, Shunji He, Guangqin Wang, Zhiyong Liu

**Affiliations:** Institute of Neuroscience, State Key Laboratory of Neuroscience, CAS Center for Excellence in Brain Science and Intelligence Technology, Chinese Academy of Sciences, Shanghai, 200031, China; University of Chinese Academy of Sciences, Beijing, 100049, China; Shanghai Center for Brain Science and Brain-Inspired Intelligence Technology, Shanghai, 201210, China

**Keywords:** Pillar cells, Deiters’ cells, inner ear, cochlear, regeneration

## Abstract

Mammalian cochlear outer hair cells (OHCs) are essential for hearing. OHC degeneration causes severe hearing impairment. Previous attempts of regenerating new OHCs from cochlear supporting cells (SCs) had yielded cells lacking Prestin, a key motor protein for OHC function. Thus, regeneration of Prestin+ OHCs remains a challenge for repairing OHC damage *in vivo*. Here, we reported that successful *in vivo* conversion of adult cochlear SCs into Prestin+ OHC-like cells could be achieved by simultaneous expression of Atoh1 and Ikzf2, two key transcriptional factors necessary for OHC development. New OHC-like cells exhibited upregulation of hundreds of OHC genes and downregulation of SC genes. Single cell transcriptomic analysis demonstrated that the differentiation status of these OHC-like cells was much more advanced than previously achieved. Thus, we have established an efficient approach to promote regeneration of Prestin+ OHCs and paved the way for repairing damaged cochlea *in vivo* via transdifferentiation of SCs.

## INTRODUCTION

Hair cells (HCs) are mammalian sound receptors that are distributed in the auditory epithelium, referred to as the organ of Corti (OC) (Wu and Kelley, 2012). Located near HCs are several supporting cell (SC) subtypes, which, from the medial side to lateral side, are Pillar cells (PCs) and Deiters’ cells (DCs) (Montcouquiol and Kelley, 2020). Auditory HCs comprise two subtypes, the inner and outer HCs (IHCs and OHCs). OHCs specifically express a unique motor protein, Prestin, encoded by *Slc26a5* (Zheng et al., 2000). Prestin-mediated electromotility enables OHCs to function as sound amplifiers, which is critical for sound detection, and *Prestin^-/-^* mice show severe hearing impairment (Liberman et al., 2002). Different from OHCs, IHCs are primary sensory cells and are innervated by type-I cochlear spiral ganglion neurons (SGNs). IHCs specifically express vGlut3, encoded by *Slc17a8*, which is required for sound information transition from IHCs to SGNs (Li et al., 2018); consequently, *vGlut3^-/-^* mice are completely deaf (Ruel et al., 2008; Seal et al., 2008). IHCs and OHCs are considered to share the same Atoh1+ progenitors (Groves et al., 2013; Tateya et al., 2019).

Atoh1 is a bHLH transcriptional factor (TF) that is necessary for specifying a general HC fate, and, accordingly, both IHCs and OHCs are lost in *Atoh1^-/-^* mice (Bermingham et al., 1999). Two additional TFs, encoded by *Insm1* and *Ikzf2*, are necessary for specifying the OHC fate or repressing the IHC fate (Chessum et al., 2018; Wiwatpanit et al., 2018). OHCs tend to transdifferentiate into IHCs in *Insm1^-/-^* and *Ikzf2* point-mutant mice. Whereas *Insm1* is only transiently expressed in differentiating OHCs (Lorenzen et al., 2015), *Ikzf2* expression is permanently maintained in differentiating and mature OHCs (Chessum et al., 2018). Unlike IHCs, OHCs are highly vulnerable to ototoxic drugs, noise, and aging. Nonmammal vertebrates such as fish and chicken can regenerate HCs from neighboring SCs in which key HC developmental genes (e.g. *Atoh1*) are reactivated (Atkinson et al., 2015), whereas mammals have lost this regenerative capacity (Janesick and Heller, 2019). PCs and DCs are physically close to OHCs, and, notably, OHCs, PCs, and DCs might share the same progenitors located in the lateral side of the OC, according to the results of recent single-cell transcriptomic analyses of cochlear cells (Kolla et al., 2020). Therefore, PCs and DCs, particularly these *Lgr5+* populations, are regarded as a favorable source for regenerating OHCs (Chai et al., 2012; McLean et al., 2017). We have shown that *in vivo* ectopic *Atoh1* can convert neonatal and juvenile SCs (primarily PCs and DCs) into nascent HCs that express early HC markers such as Myo6 and Myo7a (Liu et al., 2012a). By contrast, adult PCs and DCs are not sensitive to ectopic *Atoh1* expression *in vivo* (Kelly et al., 2012; Liu et al., 2012a), unless additional manipulations are performed (Walters et al., 2017). Nonetheless, none of the new HCs reported in previous *in vivo* studies express Prestin (Chai et al., 2012; Liu et al., 2012a; Walters et al., 2017). Therefore, it is critical to investigate how Prestin+ OHCs can be regenerated from SCs, particularly from adult SCs, in the damaged cochlea. Because ectopic *Ikzf2* in IHCs causes ectopic Prestin expression (Chessum et al., 2018), we hypothesized that *Atoh1* and *Ikzf2* would not only synergistically reprogram adult PCs and DCs into HCs, but also produce new Prestin+ OHCs.

Here, we constructed compound genetic models that allowed us to conditionally and simultaneously induce ectopic expression of *Atoh1* and *Ikzf2* in adult SCs (primarily PCs and DCs), with or without pre-damaging wild-type OHCs. Briefly, our hypothesis was supported by the results of our comprehensive genetic, transcriptomic, immunostaining, morphological and fate-mapping analyses. New Prestin+ OHC-like cells were frequently observed across the entire cochlear duct in the case of pre-damaged OHCs. To the best of our knowledge, this is the first report of the *in vivo* generation of Prestin+ OHC-like cells from adult SCs through concurrent expression of *Atoh1* and *Ikzf2*. Our findings have identified *Atoh1* and *Ikzf2* as potential targets for regenerating OHCs in hearing-impaired patients in the clinic.

## RESULTS

### Generation of conditional *Ikzf2*-overexpression genetic mouse model

Our aim was to turn on ectopic expression of *Atoh1* and *Ikzf2* specifically in adult cochlear PCs and DCs. A transgenic conditional *Atoh1*-overexpression strain, *CAG-Loxp-stop-Loxp-Atoh1*2xHA*+ (briefly, *CAG-LSL-Atoh1+*), in which 2× HA fragments were tagged at the C-terminus of Atoh1, can efficiently drive high Atoh1 expression, as we reported previously (Liu et al., 2012a). However, a similar conditional expression model for *Ikzf2* was not available. Thus, by using the CRISPR/Cas9 gene-targeting approach (Li et al., 2020a), we first generated the *Rosa26-CAG-Loxp-stop-Loxp-Ikzf2*3xHA-T2A-Tdtomato/+* knockin mouse model (briefly, *Rosa26-CAG-LSL-Ikzf2/+*), in which Ikzf2 was tagged with 3×HA fragments at its C-terminus (Figure 1-figure supplement 1A-C). Southern blotting results showed that there was no random insertion of donor DNA in the mouse genome (Figure 1-figure supplement 1D-E), and PCR-genotyping of tail DNA allowed us to readily distinguish between littermates with wild-type, heterozygous, and homozygous genotypes (Figure 1-figure supplement 1F). Heterozygous and homozygous *Rosa26-CAG-LSL-Ikzf2* mice were healthy and fertile, and did not display any apparent phenotypes. Upon Cre-mediated recombination, Tdtomato and Ikzf2 were initially translated together from the same polycistronic mRNA, and subsequently separated by the 2A peptide (Li et al., 2020b). Once triggered, the expression of Ikzf2 and Tdtomato was permanent, which enabled genetic fate-mapping analysis at single-cell resolution. Therefore, we could readily distinguish potential new OHCs (Tdtomato+) derived from adult cochlear SCs (primarily PCs and DCs in this study) from wild-type OHCs (Tdtomato-). Because high-quality commercial antibodies against neither Atoh1 nor Ikzf2 were available for immunostaining, we used an anti-HA antibody to visualize ectopic Atoh1 and Ikzf2 proteins in *CAG-LSL-Atoh1+* and *Rosa26-CAG-LSL-Ikzf2/+* strains, respectively.

**Figure 1.**
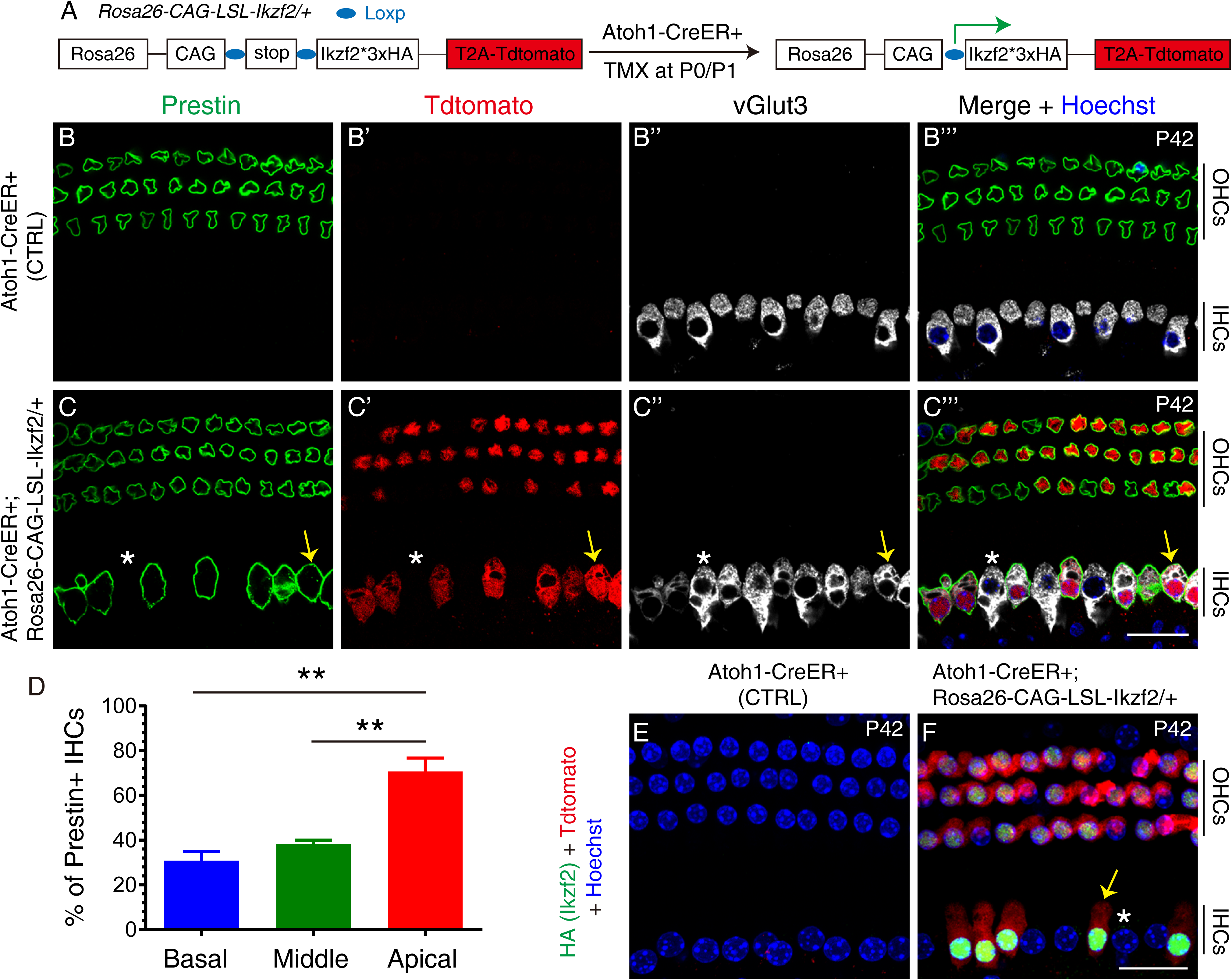
Prestin was expressed in IHCs when ectopic Ikzf2 was turned on. **(A)** Illustration of how Ikzf2 expression was turned on in HCs (both IHCs and OHCs). Tdtomato and Ikzf2 (tagged with HA) were tightly paired. *Atoh1-CreER+* is an efficient HC cre driver at neonatal ages. **(B-C’’’)** Triple labeling for Prestin, Tdtomato, and vGlut3 in P42 cochlear samples: control *Atoh1-CreER+* (B-B’’’) and experimental *Atoh1-CreER+; Rosa26-CAG-LSL-Ikzf2/+* (C-C’’’). Prestin was only expressed in wild-type OHCs (B-B’’’). Arrows in (C-C’’’): Tdtomato+/vGlut3+/Prestin+ IHC; asterisks in (C-C’’’): vGlut3+/Tdtomato-IHC that did not turn on Prestin expression in experimental mice. **(D)** Quantification of Prestin+ IHCs. More Prestin+ IHCs were present in the apical turn than in basal and middle turns. ** p < 0.01. **(E-F)** Co-staining of HA (Ikzf2) and Tdtomato in control (E) and experimental (F) mice at P42. HA+/Tdtomato+ cells were present in experimental mice only. Arrow/asterisk in (F): IHC with/without HA (Ikzf2) expression. All Tdtomato+ cells were HA+ (Ikzf2-expressing) cells, and vice versa. Scale bars: 20 μm.

### Ectopic Ikzf2 was able to induce Prestin expression in IHCs

Before testing *Ikzf2* expression here as a method to regenerate OHCs from adult cochlear SCs, we determined whether functional *Ikzf2* was efficiently induced in *Rosa26-CAG-LSL-Ikzf2/+* mice. Prestin is expressed in OHCs exclusively in wild type mice (Fang et al., 2012; Liberman et al., 2002; Zheng et al., 2000), but it is ectopically turned on in IHCs infected with an Anc80-virus expressing *Ikzf2* (Chessum et al., 2018). Therefore, the criterion we used was that Prestin expression must be triggered in IHCs upon crossing *Rosa26-CAG-LSL-Ikzf2/+* with the transgenic strain *Atoh1-CreER+*, in which Cre activity is restricted to HCs (both IHCs and OHCs) (Chow et al., 2006; Cox et al., 2012). Both *Atoh1-CreER+* (control group) and *Atoh1-CreER+; Rosa26-CAG-LSL-Ikzf2/+* (experimental group) mice were administered tamoxifen at postnatal day 0 (P0) and P1, and analyzed at P42 (Figure 1A and Figure 1-figure supplement 2). We also comprehensively analyzed another control strain, *Rosa26-CAG-LSL-Ikzf2/+*, and we did not detect either HA+ (Ikzf2+) or Tdtomato+ cells in the cochlea in this strain, which was not described further here.

We primarily focused on Prestin, HA (Ikzf2), and Tdtomato expression patterns in IHCs, although Prestin+ OHCs were also targeted. Neither Prestin nor Tdtomato was expressed in IHCs of control mice (n = 3) at P42 (Figure 1B-B’’’). By contrast, Prestin was ectopically expressed in IHCs that were vGlut3+/Tdtomato+ (arrows in Figure 1C-C’’’) but not IHCs that were vGlut3+/Tdtomato-(asterisks in Figure 1C-C’’’) in the experimental mice at P42 (n = 3). All Tdtomato+ IHCs were Prestin+ and all Prestin+ IHCs were Tdtomato+. Quantification revealed that 30.8% ± 4.2%, 38.4% ± 1.6%, and 70.7% ± 6.0% of IHCs were Prestin+ in basal, middle, and apical turns, respectively, at P42 (Figure 1D and Figure 1-figure supplement 2); thus, the apical turn harbored more Prestin+ IHCs than the basal and middle turns, in accord with the higher Cre efficiency of *Atoh1-CreER+* in the apex (Cox et al., 2012). We also confirmed that all Tdtomato+ (Prestin+) IHCs expressed HA (Ikzf2) and vice versa (Figure 1E and F), which validated the co-expression of Ikzf2 and Tdtomato. This result also suggested that Ikzf2 derepressed Prestin expression in IHCs in a cell-autonomous manner. Moreover, Tdtomato+ OHCs were expected to express both endogenous and ectopic Ikzf2, but the OHCs appeared normal, which suggests that these cells can tolerate additional Ikzf2 expression until at least P42 (Figure 1C-C’’’ and Figure 1F).

Together, our results showed that the expression level of Ikzf2 derived from the *Rosa26-CAG-LSL-Ikzf2/+* strain was sufficiently high to drive Prestin expression in IHCs. These Tdtomato+/Ikzf2+/Prestin+ IHCs were expected to be distinct from wild-type IHCs, because these cells might partially transdifferentiate into OHCs or at least express OHC genes (Chessum et al., 2018). To summarize, we confirmed that *Rosa26-CAG-LSL-Ikzf2/+* was a powerful genetic model suitable for inducing functional Ikzf2 expression in adult cochlear SCs.

### Ikzf2 alone failed to convert adult PCs and DCs into HCs

We next determined whether *Ikzf2* alone can reprogram adult cochlear SCs into Prestin+ OHCs. No Tdtomato+ HCs (neither IHCs nor OHCs) were observed in *Fgfr3-iCreER+; Ai9/+* (briefly, *Fgfr3-Ai9*) mice when tamoxifen was administered at P30/P31 and the analysis was performed at P60 (arrows in Figure 2A-A’’’). However, most SCs (primarily PCs and DCs) were Tdtomato+ (inset in Figure 2A’), which agreed with our previous reports (Liu et al., 2012a; Liu et al., 2012b). We also confirmed that Atoh1 alone failed to convert adult cochlear SCs into HCs in the *Fgfr3-iCreER+; CAG-LSL-Atoh1+* model (briefly, *Fgfr3-Atoh1*) (Figure 2B-B’’’), same as we reported previously (Liu et al., 2012a).

**Figure 2.**
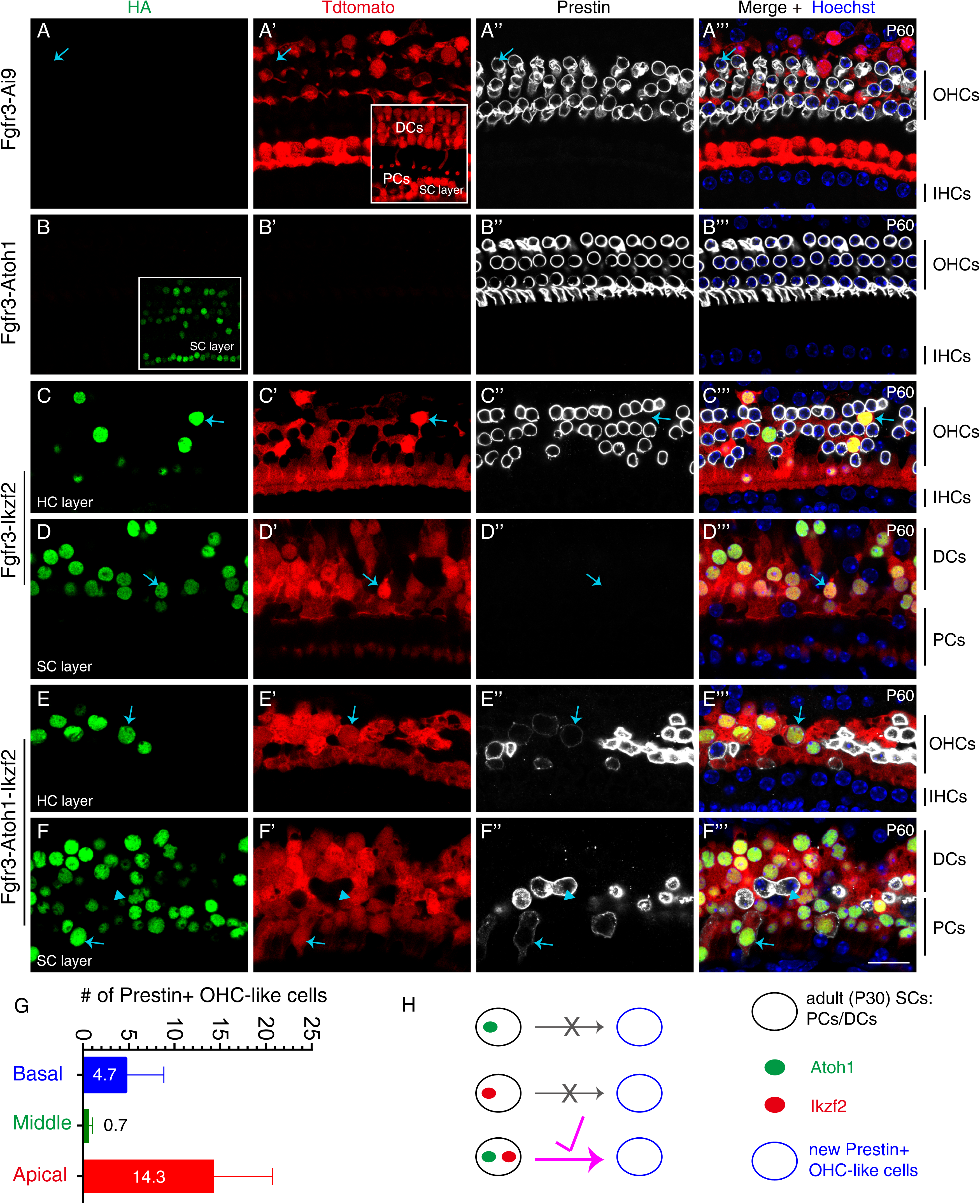
Atoh1 and Ikzf2 together converted adult PCs and DCs into OHC-like cells at low efficiency. Triple labeling for HA, Tdtomato, and Prestin in four different mouse genetic models that were administered tamoxifen at P30 and P31 and then analyzed at P60. **(A-A’’’)** All Tdtomato+ were SCs (primarily PCs and DCs) in *Fgfr3-iCreER+; Ai9/+ (Fgfr3-Ai9)* mice. Inset in (A’): confocal image scanned at SC layer. Arrows in (A-A’’’): Prestin+/Tdtomato-OHC. **(B-B’’’)** No Tdtomato signal was detected in *Fgfr3-iCreER+; CAG-LSL-Atoh1+ (Fgfr3-Atoh1)* mice, and no HA+/Prestin+ cells were observed. Inset in (B): confocal image scanned at SC layer. **(C-D’’’)** Confocal images scanned at HC layer (C-C’’’) and SC layer (D-D’’’) in cochlear samples from *Fgfr3-iCreER+; Rosa26-CAG-LSL-Ikzf2/+* (*Fgfr3-Ikzf2*) mice. Arrows in both layers: two cells that were HA+/Tdtomato+ but did not express Prestin. Alignment of wild-type Prestin+ OHCs was abnormal. **(E-F’’’)** Confocal images scanned at HC layer (E-E’’’) and SC layer (F-F’’’) in cochlear samples from *Fgfr3-iCreER+; CAG-LSL-Atoh1+; Rosa26-CAG-LSL-Ikzf2/+* (*Fgfr3-Atoh1-Ikzf2*) mice. According to different cellular morphologies and location, arrows in (E-E’’’) indicate an OHC-like cell that was HA+/Tdtomato+/Prestin+ and derived from adult DCs; by contrast, the arrows in (F-F’’’) indicate another OHC-like cell derived from adult PCs. Arrowheads: Prestin+/Tdtomato-wild-type OHC appearing in SC layer. Prestin expression in the new OHC-like cells was lower than that in wild-type OHCs. **(G)** Quantification of OHC-like cells throughout entire cochlear turns in the *Fgfr3-Atoh1-Ikzf*2 model. Data are presented as means ± SEM (n=3). OHC-like cells were reproducibly observed, but the cell numbers were low and showed large variations. **(H)** Summary of reprogramming outcomes in the three models studied here. OHC-like cells were present only when *Atoh1* and *Ikzf2* were concurrently reactivated in adult cochlear SCs. Scale bars: 20 μm.

As expected, numerous Tdtomato+ cells were observed in cochleae of *Fgfr3-iCreER+; Rosa26-CAG-LSL-Ikzf2/+* (briefly, *Fgfr3-Ikzf2*) mice that were given tamoxifen at P30/P31 and analyzed at P60, in both the HC layer (Figure 2C-C’’’) and the SC layer (Figure 2D-D’’’). Again, all Tdtomato+ cells expressed HA and vice versa (arrows in Figure 2C-D’’’). However, none of these Tdtomato+/HA+ cells expressed Prestin (arrows in Figure 2C-D’’’). Furthermore, we also did not detect any Tdtomato+/Myo6+ cells. We noted the loss of endogenous OHCs (Prestin+/Tdtomato-) throughout the cochlear duct, particularly at the basal turn, which was likely a secondary effect of ectopic Ikzf2 expression in adult cochlear SCs. Collectively, these results suggested that Ikzf2 alone was not sufficient for converting adult cochlear SCs into nascent Myo6+ HCs or Prestin+ OHCs. Thus, in terms of cell-fate conversion, more barriers might exist between adult cochlear SCs and OHCs than between IHCs and OHCs.

### Ikzf2 and Atoh1 together converted, at low efficiency, adult PCs and DCs into Prestin+ OHC-like cells

Considering the synergistic effects reported among multiple TFs such as Six1, Atoh1, Pou4f3, and Gata3 and between Myc and Notch signaling (Costa et al., 2015; Menendez et al., 2020; Shu et al., 2019; Walters et al., 2017), we hypothesized that concomitant induction of *Ikzf2* and *Atoh1* might convert adult cochlear SCs into OHCs. We tested this by analyzing *Fgfr3-iCreER+; CAG-LSL-Atoh1+; Rosa26-CAG-LSL-Ikzf2/+* mice (briefly, *Fgfr3-Atoh1-Ikzf2*) that were given tamoxifen at P30/P31 and analyzed at P60. Again, Tdtomato+ SCs were abundant within the OC, and 88.0% ± 2.7%, 94.1% ± 4.3%, and 98.2% ± 1.8% of HA+ cells were Tdtomato+ in the basal, middle, apical turns, respectively (n = 3). The finding that most of the Tdtomato+ cells were HA+ further confirmed the high Cre activity in the *Fgfr3-iCreER* model. The small fraction of HA+/Tdtomato-cells represented populations in which ectopic Atoh1 but not Ikzf2 was expressed due to independent Cre-recombination events in the two loci. Conversely, no Tdtomato+/HA-cells were observed because Tdtomato and HA are tightly paired in the *Rosa26-CAG-LSL-Ikzf2/+* model.

Unlike in the case of *Ikzf2* induction alone (Figure 2C-D’’’), Tdtomato+/Prestin+ cells were occasionally observed in *Fgfr3-Atoh1-Ikzf2* mice (n = 3) at P60 (arrows in Figure 2E-F’’’). These cells were defined as new OHC-like cells in our study because they were derived from the original Tdtomato+ cochlear SCs (PCs and DCs) but were not identical to wild-type adult OHCs yet. These OHC-like cells were distributed in both the HC layer (Figure 2E-E’’’) and the SC layer (Figure 2F-F’’’). However, the numbers of new OHC-like cells detected were only 4.7 ± 4.2, 0.7 ± 0.3, and 14.3 ± 6.4 throughout the entire basal, middle, and apical turns, respectively (Figure 2G). Moreover, Prestin protein expression was substantially lower in these OHC-like cells (arrows in Figure 2E-F’’’) than in wild-type endogenous OHCs, which expressed Prestin but not Tdtomato (arrowheads in Figure 2F-F’’’). Again, endogenous OHC loss occurred in all cochlear turns (Figure 2E-F’’’). Collectively, these results supported the conclusion that *Atoh1* and *Ikzf2* together reprogrammed, albeit at low efficiency, adult cochlear SCs into Prestin+ OHC-like cells, but neither gene alone triggered this conversion (Figure 2H). Therefore, we next sought to test whether pre-damaging OHCs would boost the reprogramming efficiency and generate increased numbers of Prestin+ OHC-like cells.

### Generation of *Prestin-DTR/+* model for OHC damage

To damage adult OHCs *in vivo*, we used genetic and pharmacological approaches. The diphtheria toxin (DT)/DT receptor (DTR) system has been successfully used to damage HCs in the inner ear (Cox et al., 2014; Golub et al., 2012; Tong et al., 2015). Thus, we generated a new knockin mouse model, *Prestin-P2A-DTR/+* (briefly, *Prestin-DTR/+*), in which the P2A-DTR fragment was inserted immediately before the stop codon (*TAA*) of *Prestin* (Figure 3-figure supplement 1A-C). DTR expression was entirely controlled by the endogenous *Prestin* promoter and/or enhancers and restricted to OHCs, and Prestin expression itself was intact. Southern blotting results confirmed the absence of random insertion of donor DNA in the genome (Figure 3-figure supplement 1D and E), and tail-DNA PCR allowed us to readily distinguish between wild-type *Prestin* and post-gene-targeting alleles (Figure 3-figure supplement 1F).

In the absence of DT treatment, co-staining for the OHC-marker Prestin and IHC-marker vGlut3 revealed that OHCs were normal in *Prestin-DTR/+* mice (n = 3) at P42 (Figure 3A and A’). By contrast, after a single injection of DT (20 ng/g, body weight) at P36, severe OHC loss was observed in *Prestin-DTR/+* mice (n = 3) at P42, and only very few OHCs were sporadically detected throughout the cochlear duct (arrowhead in Figure 3B’). The debris of dying OHCs were frequently observed at P42 (arrows in Figure 3B’) but disappeared by P60 (further described below; see Figure 4). Conversely, IHCs appeared normal at P42 under DT treatment, which confirmed that DTR was specifically expressed in OHC and absent in IHCs (Figure 3B and B’). Furthermore, the results of auditory brainstem response (ABR) measurement demonstrated that the thresholds at distinct frequencies in *Prestin-DTR/+* mice (n=3) treated with DT were significantly higher than those in control *Prestin-DTR/+* mice (n=3) not treated DT (Figure 3C). Accordingly, the ratio of OHC numbers to IHC number was ∼3.23 in control (blue in Figure 3D), but was significantly decreased, to ∼0.19, in *Prestin-DTR/+* mice treated with DT (red in Figure 3D). Together, these results showed that DT treatment caused, within 6 days, marked hearing impairment due to OHC loss, and further indicated that *Prestin-DTR/+* represented a powerful mouse model for specifically damaging wild-type OHCs.

**Figure 3.**
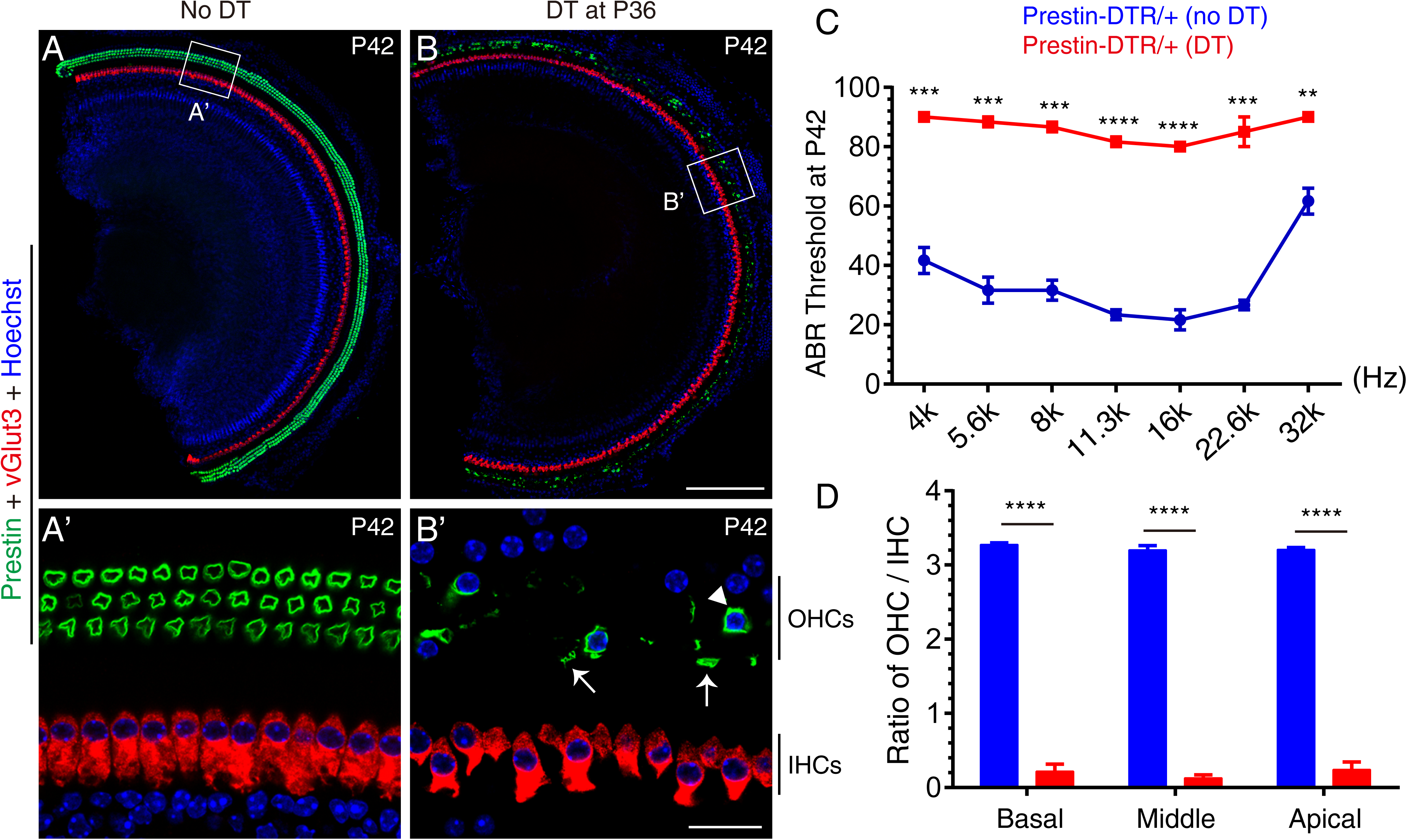
Damaging adult OHCs specifically by using genetic and pharmacological approaches. (A-B’) *Prestin-DTR/+* mice were treated without (A-A’, control) or with (B-B’, experimental) diphtheria toxin (DT) at P36 and analyzed at P42. Samples were co-stained for Prestin and vGlut3. (A’) and (B’): magnified images of indicated square areas in (A) and (B). DT treatment led to rapid OHC death within 6 days and only a few OHCs remained (arrowhead in B’), and debris of dying OHCs were frequently detected (arrows in B’). Much of the green signal in (B) was from the debris of dying OHCs. **(C)** Auditory brainstem response (ABR) measurement of *Prestin-DTR/+* mice treated without (blue line) or with (red line) DT. ABR thresholds were significantly higher throughout cochlear ducts after DT treatment. **(D)** Ratios of OHCs to IHCs in control mice (blue) and experimental mice (red) in the same confocal scanning areas; OHC numbers were significantly decreased in the experimental mice. ** p<0.01, *** p<0.001, **** p<0.0001. Scale bars: 200 μm (B), 20 μm (B’).

**Figure 4.**
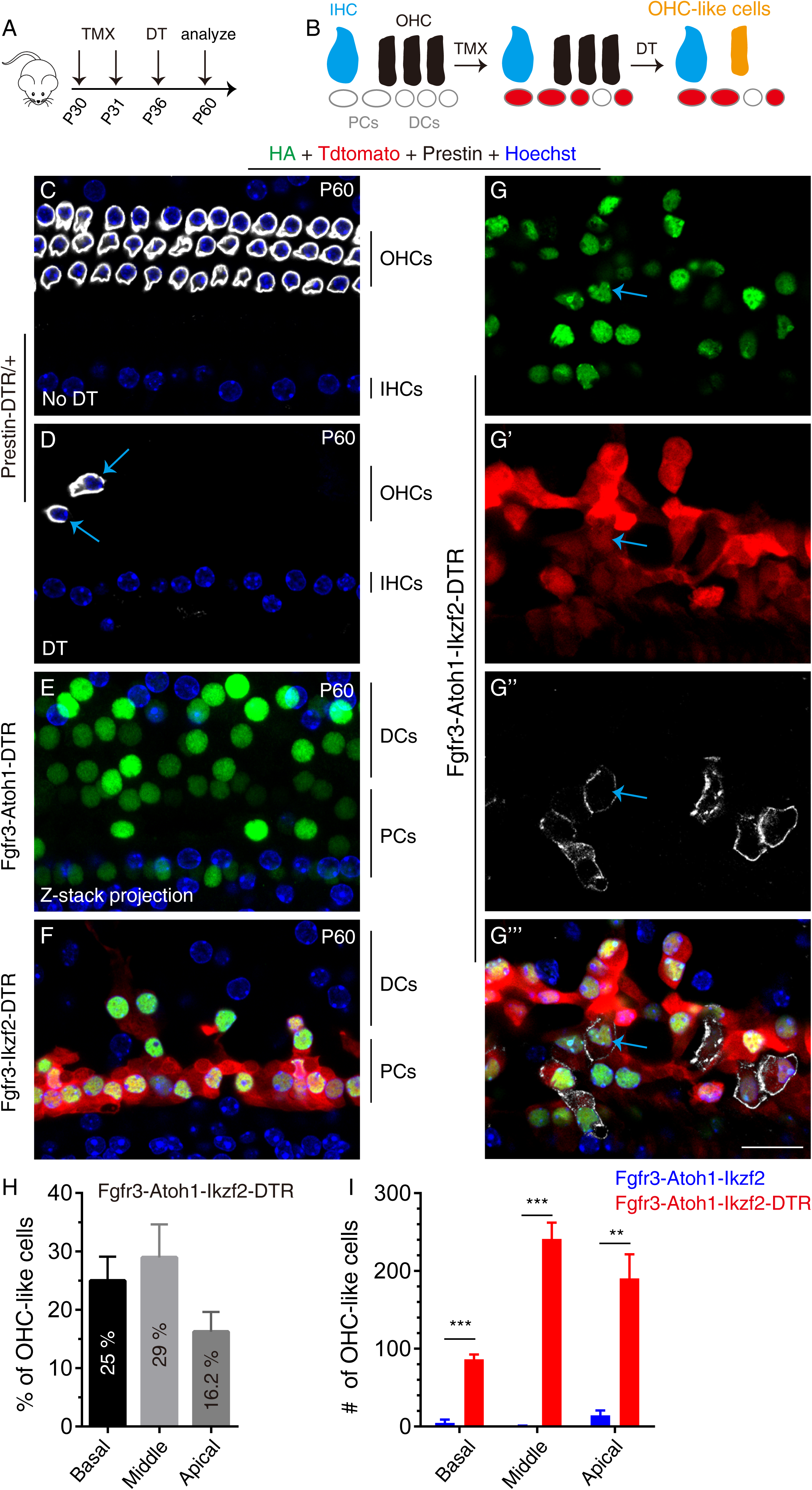
Damaging wild-type OHCs enhanced reprogramming efficiency of Atoh1 and Ikzf2 in adult SCs. **(A)** Identical treatments were applied to the distinct genetic mouse models: tamoxifen treatment at P30 and P31, followed by DT treatment at P36 and analysis at P60. **(B)** Drawing illustrating key events at the cellular level: turning on *Atoh1* and *Ikzf2* in adult PCs and DCs that were also permanently labeled with Tdtomato for the subsequent fate-mapping analysis. **(C-G’’’)** Triple labeling for HA, Tdtomato, and Prestin in four models: (1) *Prestin-DTR/+* (C and D), (2) *Fgfr3-iCreER+; CAG-LSL-Atoh1+; Prestin-DTR/+* (*Fgfr3-Atoh1-DTR*; E), (3) *Fgfr3-iCreER+; Rosa26-CAG-LSL-Ikzf2/+; Prestin-DTR/+* (*Fgfr3-Ikzf2-DTR*; F), and (4) *Fgfr3-iCreER+; CAG-LSL-Atoh1+; Rosa26-CAG-LSL-Ikzf2/+; Prestin-DTR/+*(*Fgfr3-Atoh1-Ikzf2-DTR*; G-G’’’’). Relative to *Prestin-DTR/+* mice not treated with DT (C), DT-treated *Prestin-DTR/+* mice harbored very few normal Prestin+ OHCs at P60 (arrows in D). Debris of dying OHCs had disappeared. No OHC-like cells were observed in the first three models, but Tdtomato+/HA+/Prestin+ OHC-like cells (arrows in G-G’’’) were present in the *Fgfr3-Atoh1-Ikzf2-DTR* model. **(H)** Percentages of OHC-like cells at different cochlear turns in *Fgfr3-Atoh1-Ikzf2-DTR* mice. **(I)** Comparison of OHC-like cell numbers between *Fgfr3-Atoh1-Ikzf2-DTR* and *Fgfr3-Atoh1-Ikzf2* models (without damaging adult wild-type OHCs). *Fgfr3-Atoh1-Ikzf2-DTR* mice harbored considerably more OHC-like cells than *Fgfr3-Atoh1-Ikzf2* mice. ** p<0.01, *** p<0.001. Scale bars: 20 μm.

### Conversion of adult cochlear SCs into Prestin+ OHC-like cells by Ikzf2 and Atoh1 was considerably enhanced when endogenous OHCs were damaged

In nonmammalian vertebrates, HC loss triggers the regeneration of HCs through a cell-fate change in SCs (Janesick and Heller, 2019; Stone and Cotanche, 2007; Warchol and Corwin, 1996). Therefore, we tested whether the damage of endogenous wild-type adult OHCs coupled with the ectopic expression of Ikzf2 and Atoh1 in adult cochlear SCs would efficiently generate OHC-like cells; for this, we used four genetic models: (1) Prestin-DTR/+; (2) Fgfr3-iCreER+; CAG-LSL-Atoh1+; Prestin-DTR/+ (Fgfr3-Atoh1-DTR); (3) Fgfr3-iCreER+; Rosa26-CAG-LSL-Ikzf2/+; Prestin-DTR/+ (Fgfr3-Ikzf2-DTR); and (4) Fgfr3-iCreER+; CAG-LSL-Atoh1+; Rosa26-CAG-LSL-Ikzf2/+; Prestin-DTR/+ (Fgfr3-Atoh1-Ikzf2-DTR). We first turned on dual expression of Atoh1 or Ikzf2 or both (by injecting tamoxifen at P30 and P31) and then triggered OHC damage 6 days later (by DT treatment at P36) (Figure 4A and B). This strategy allowed us to precisely and permanently label adult SCs (mainly PCs and DCs) with HA and Tdtomato before OHC damage and thus facilitated the subsequent fate-mapping analysis for determining whether Prestin+ OHC-like cells were produced. The reverse order of the experimental procedure was not used so as to avoid the possibility of OHC damage leading to changes in the Cre-expression pattern in *Fgfr3-iCreER+* mice.

Unlike in control *Prestin-DTR/+* mice not treated with DT, in which three well-aligned rows of Prestin+ OHCs were observed at P60 (Figure 4C), only a few OHCs were occasionally detected in *Prestin-DTR/+* mice treated DT (arrows in Figure 4D). Moreover, in contrast to the case at P42 (arrows in Figure 3B’), we detected no debris of dying OHCs at P60 (Figure 4D). Furthermore, no HA+/Prestin+ cells were identified in either *Fgfr3-Atoh1-DTR* mice (Figure 4E) or *Fgfr3-Ikzf2-DTR* mice (Figure 4F) at P60, which suggested that damaging wild-type OHCs did not promote production of the Prestin+ new OHC-like cells when Atoh1 or Ikzf2 was overexpressed alone in adult cochlear SCs.

Conversely, Tdtomato+/HA+/Prestin+ OHC-like cells were frequently observed in *Fgfr3-Atoh1-Ikzf2-DTR* mice at P60. Confocal scanning of the entire cochlear duct and quantification (n = 3) revealed the presence of 359.3 ± 46.2, 878.0 ± 118.7, and 1195 ± 81.6 Tdtomato+/HA+ cells in the basal, middle, and apical turns, respectively, among which 86.3 ± 6.3, 241.0 ± 21.1, and 190.3 ± 31.1 cells were the new OHC-like cells (arrows in Figure 4G-G’’’). By normalizing the numbers of OHC-like cells against the total number of Tdtomato+/HA+ cells per sample, we found that 25.0% ± 4.1%, 29.0% ± 5.7%, and 16.2% ± 3.4% of adult cochlear SCs expressing both Ikzf2 and Atoh1 transformed into OHC-like cells in the basal, middle, and apical turns, respectively (Figure 4H). Comparison with the results obtained for *Fgfr3-Atoh1-Ikzf2* mice at P60 (Figure 2E-G) revealed not only a significant increase in the number of Prestin+ OHC-like cells at each cochlear turn in *Fgfr3-Atoh1-Ikzf2-DTR* mice (Figure 4I), but also a general elevation of Prestin expression in individual cells (Figure 4G-G’’’). However, the Prestin levels in these OHC-like cells were considerably lower than those in wild-type OHCs (Figure 4C vs Figure 4G’’). Together, our results demonstrated that damaging endogenous OHCs markedly enhanced the reprogramming efficiency of Ikzf2 and Atoh1, and thus considerably increased numbers of OHC-like cells derived from adult cochlear SCs.

### Initial cell-fate transition from SCs to general HCs was followed by a second switch from general HCs to OHC-like cells

We next determined when these adult cochlear SC-derived OHC-like cells or nascent new HCs emerged (Figure 4-figure supplement 1A-C’’’). Nascent new HCs were Tdtomato+/Myo6+ but had not turned on Prestin expression yet (arrowheads in Figure 4-figure supplement 1B-C’’’); P42 was the earliest age at which the nascent new HCs were detected, and Myo6 expression was weak at this stage. Scanning of the entire cochlear duct at P42 and quantification revealed the presence of only 29.0 ± 13.5, 66.0 ± 35.23, and 52.7 ± 25.4 nascent new HCs throughout the basal, middle, and apical turns (n=3), respectively (black in Figure 4-figure supplement 1D). However, no Tdtomato+/Prestin+ OHC-like cells were detected at P42.

Four days later, at P46 (n = 4), there were 56.5 ± 28.8, 118.0 ± 61.2, and 111.0 ± 57.3 new HCs that were Tdtomato+/Myo6+ (gray in Figure 4-figure supplement 1D and E), and, of these, 27.8 ± 15.0, 43.5 ± 25.3, and 36.8 ± 21.0 were Tdtomato+/Myo6+/Prestin+ (OHC-like cells; green in Figure 4-figure supplement 1E); thus, OHC-like cells accounted for 49.2% (27.8/56.5), 36.9% (43.5/118), and 33.2% (36.8/111) of total new HCs in basal, middle, and apical turns, respectively. The remaining Tdtomato+/Myo6+/Prestin-cells were defined as nascent HCs, and the Tdtomato+/Myo6-/Prestin-cells were defined as SCs that failed to become HCs. Here, we sorted new HCs into nascent HCs and OHC-like cells based solely on absence or presence of Prestin. Among the 4 mice analyzed, 2 mice harbored substantially fewer Tdtomato+/Myo6+ cells, which caused the large variations in numbers; nevertheless, the overall trend was that the higher the number of Tdtomato+/Myo6+ cells observed, the higher the number of OHC-like cells observed. Notably, we did not detect even a single Tdtomato+/Prestin+ cell not expressing Myo6. Together, these findings suggested that the generation of new OHC-like cells generally involved an initial cell-fate transition from SCs (PCs and DCs) into nascent HCs by P42 (12 days after turning on *Atoh1* and *Ikzf2*, and 6 days after DT treatment), and this was followed by a second transition from nascent HCs into OHC-like cells.

### Hair bundles were present in the new OHC-like cells

Scanning electron microscopy (SEM) was used to examine the hair bundles (or stereocilia) of the Prestin+ new OHC-like cells at P60 (Figure 5). The staircase-shaped hair bundles are the sites where mechanoelectrical transduction (MET) channels are distributed and are critical for hearing (Corey and Holt, 2016; Douguet and Honore, 2019; Wu and Muller, 2016). The regular V- or W-shaped hair bundles were present in wild-type OHCs from *Prestin-DTR/+* mice not treated with DT (control) (Figure 5A and A’), whereas very few hair bundles remained in *Prestin-DTR/+* mice treated with DT (Figure 5B). This agreed with the results of our immunostaining assays (Figure 4D). Intriguingly, we frequently detected stereocilia with a single long bundle but lacking the staircase shape in *Fgfr3-Atoh1-Ikzf2-DTR* mice at P60 (Figure 5C-C’). Stereocilia with several long bundles were seldom observed (inset in Figure 5C). These are likely the hair bundles of OHC-like cells (or nascent HCs), because such hair bundles were not observed in *Prestin-DTR/+* mice treated with or without DT (Figure 5A-B). ABR measurement results showed that the thresholds at distinct frequencies were markedly higher in *Fgfr3-Atoh1-Ikzf2-DTR* mice (n=6, red line in Figure 5D) than in *Prestin-DTR/+* mice without DT treatment (n=3, black line in Figure 5D). However, no significant hearing improvement (lowering of threshold) was recorded between *Fgfr3-Atoh1-Ikzf2-DTR* and *Prestin-DTR/+* mice upon DT treatment (n=5, blue line in Figure 5D). Notably, loss of IHCs were observed at P60 (asterisks in Figure 5B and C). It should be a secondary effect caused by OHC death or disruption of the OC structure because no IHC death was observed at P42 (Figure 3). Collectively, these results showed that the new OHC-like cells did not yet completely resemble wild-type OHCs, at least in terms of hair-bundle structure and Prestin expression. The extent to which OHC-like cells were similar to wild-type OHCs was next addressed using single-cell RNA-Seq analysis.

**Figure 5.**
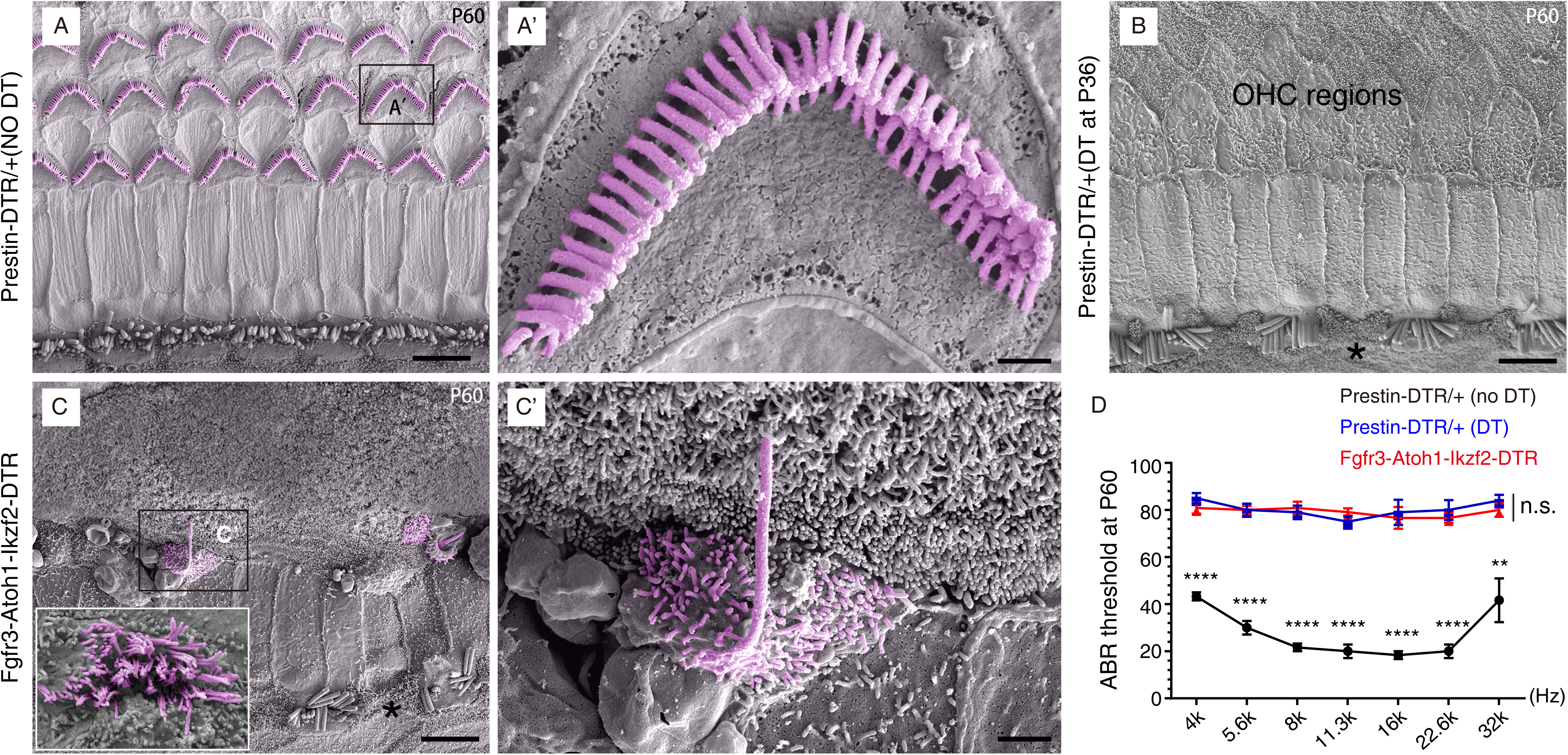
Hair bundles were present in OHC-like cells. Scanning electron microscopy (SEM) analysis of samples from three different mouse models at P60. **(A-A’)** OHCs harbored V- or W-shaped HC bundles in *Prestin-DTR/+* mice not treated with DT. (A’): high-magnification view of black rectangle in (A). **(B)** Almost all OHCs disappeared in *Prestin-DTR/+* mice upon DT treatment at P36. Black asterisk: one IHC that was absent. **(C)** Immature hair bundles were frequently detected in the *Fgfr3-iCreER+; CAG-LSL-Atoh1+; Rosa26-CAG-LSL-Ikzf2/+; Prestin-DTR/+* (*Fgfr3-Atoh1-Ikzf2-DTR*) model, but not in (A) and (B). These hair bundles were expected to originate from OHC-like cells. (C’): high-magnification view of black rectangle in (C). Inset in C: HC bundles that were observed relatively less frequently. Black asterisk: one IHC that was missing. **(D)** ABR measurements of these three models. Relative to the threshold in *Prestin-DTR/+* mice not treated with DT (black line), the ABR thresholds in *Prestin-DTR/+* treated with DT (blue line) and in *Fgfr3-Atoh1-Ikzf2-DTR* mice (red line) were significantly increased. The blue and red lines showed no statistically significant difference at any frequency (n.s.). ** p<0.01, **** p<0.0001. Scale bars: 5 μm (A, B, C), 1 μm (C’), and 500 nm (A’).

### Single-cell RNA-Seq revealed unique genes enriched in adult wild-type OHCs and SCs

To perform single-cell RNA-Seq on adult OHCs and SCs, we manually picked 17 wild-type Tdtomato+ OHCs from *Prestin-CreER/+; Ai9/+* mice at P30, and 16 wild-type Tdtomato+ SCs (primarily PCs and DCs) from *Fgfr3-iCreER+; Ai9/+* mice at P60 (Figure 6A). All cells were identified based on their endogenous Tdtomato fluorescence and were picked and washed thrice under a fluorescence microscope before final collection in PCR tubes, after which RNA-Seq libraries were prepared using the Smart-Seq approach (smartseq) (Figure 6A). Manual picking combined with smartseq has been successfully used in previous gene-profiling studies on adult cochlear HCs and SGNs (Li et al., 2020a; Liu et al., 2014; Shrestha et al., 2018). Because smartseq involves full-length cDNA sequencing, the method provides higher gene coverage than 10× genomic single-cell RNA-Seq, which only detects the 3ʹ-end of coding genes (Li et al., 2020a; Petitpre et al., 2018; Shrestha et al., 2018; Sun et al., 2018; Yamashita et al., 2018).

**Figure 6.**
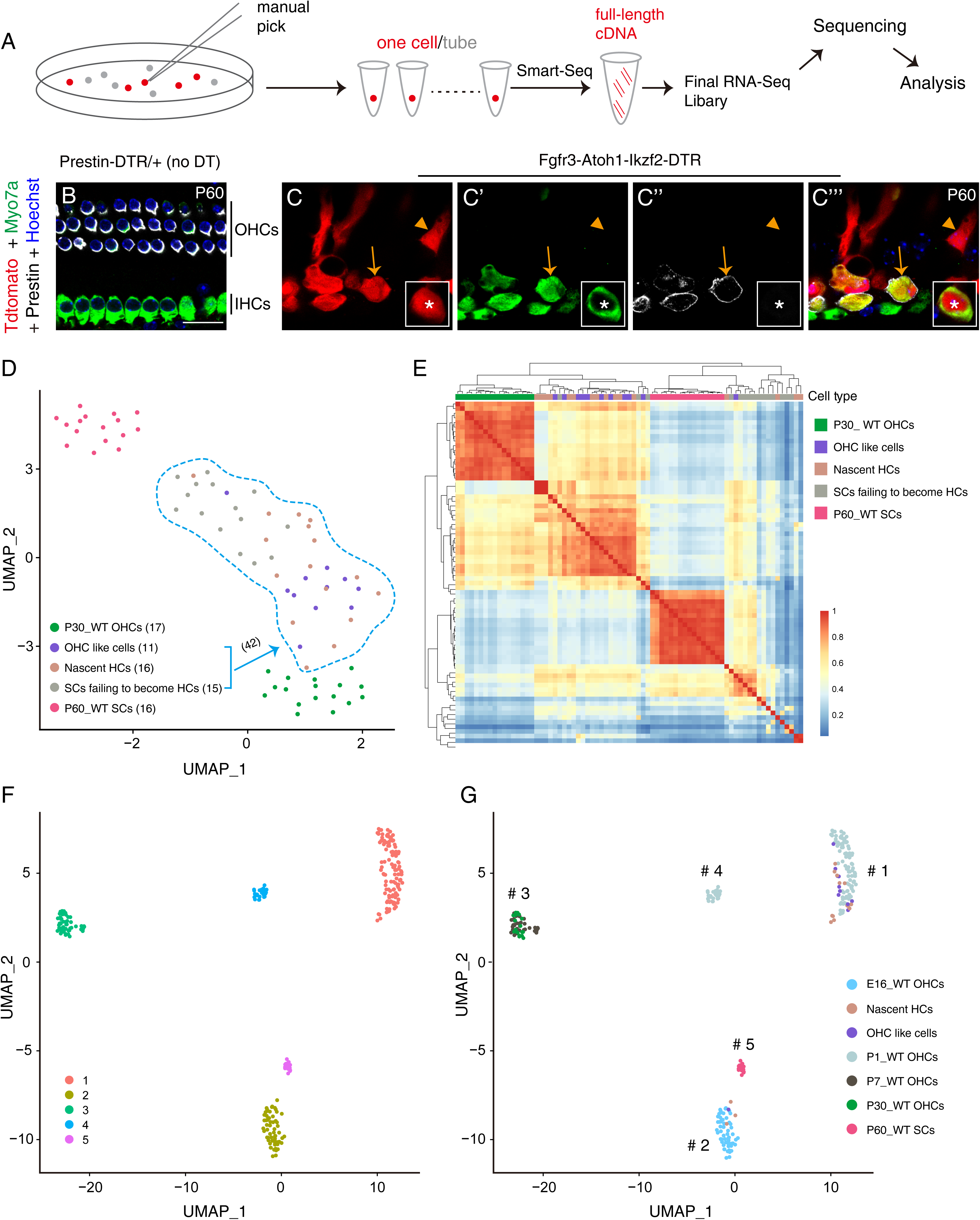
OHC-like cells were most similar to wild-type OHCs at ∼P1. **(A)** Drawing illustrating manual picking of Tdtomato+ cells (of different cell types from three different models) and single-cell RNA-Seq. **(B-C’’’)** Triple labeling for Myo7a, Prestin, and Tdtomato in cochlear samples from control (B) and *Fgfr3-Atoh1-Ikzf2-DTR* (C) mice at P60. Arrows: Tdtomato+/Myo7a+/Prestin+ OHC-like cell; arrowheads: Tdtomato+/Myo7a-/Prestin-cell, defined as SC that failed to become an HC; asterisks: nascent HC that was Tdtomato+/Myo7a+/Prestin- (inset in C-C’’’). **(D)** UMAP analysis of five different cell types together. All cells were Tdtomato+, but were picked from three different models. The 42 Tdtomato+ cells within the light-blue dotted lines were from *Fgfr3-Atoh1-Ikzf2-DTR* mice at P60; these cells were divided into three sub-clusters. **(E)** Pearson correlation-coefficient analysis of the same five cell types as in (D). **(F-G)** UMAP analysis of all cells in (D-E), except for SCs that failed to become HCs, together with wild-type OHCs at E16, P1, and P7 (E16_WT OHCs, P1_WT OHCs, and P7_WT OHCs; detailed information in Supplemental Figure 8).

We first compared gene profiles between wild-type OHCs at P30 (P30_WT OHCs) and wild-type SCs at P60 (P60_WT SCs), and we identified 1324 and 2344 genes enriched respectively in adult OHCs and adult SCs (Figure 6-figure supplement 1A); Supplemental File 1 contained the entire list of enriched genes. The OHC-enriched genes included genes that are recognized to be highly expressed in adult OHCs, such as *Myo7a*, *Ocm*, *Prestin (Slc26a5)*, *Ikzf2*, *Espn*, *Tmc1*, *Cib2*, *Lhfpl5*, *Lmo7*, *Lbh*, and *Sri* (Chessum et al., 2018; Du et al., 2019; Giese et al., 2017; Liu et al., 2014; Ranum et al., 2019; Xiong et al., 2012; Zheng et al., 2000). As expected, previously identified IHC-specific genes such as *Slc7a14* (blue arrows in Figure 6-figure supplement 1A) were either not detected or minimally detected in our picked Tdtomato+ OHCs (Liu et al., 2014); by contrast, previously known pan-SC markers, such as *Sox2* and *Sox10*, and two recently identified DC-specific genes, *Bace2* and *Ceacam16*, were included among the SC-enriched genes (Li et al., 2018; Ranum et al., 2019). Gene Ontology (GO) enrichment analysis was performed on the OHC- or SC-enriched genes (Figure 6-figure supplement 1B and C); the results confirmed that the OHC-enriched genes were involved in sensory perception of sound, inner ear morphogenesis, neurotransmitter secretion, and stereocilium organization (Figure 6-figure supplement 1B), and that the SC-enriched genes were involved in functions such as lipid metabolic process, regulation of cell shape, and actin cytoskeleton organization (Figure 6-figure supplement 1C). This agreed with the finding that PCs are critical for the formation of the tunnel of Corti featuring a unique morphology. The OHC- and SC-enriched gene lists of each GO category were summarized in Supplemental Files 2 and 3, respectively. Overall, the results showed that the picked adult wild-type OHCs and SCs were pure and that our single-cell RNA-Seq data were of high quality. Therefore, these genes specifically enriched in adult wild-type OHCs and SCs served as references in our characterization of the new HCs, particularly the OHC-like cells.

Although Myo6 protein is known to be enriched in OHCs, *Myo6* mRNA was not significantly enriched in adult OHCs, because, in our hands, the mRNA was also detected (albeit at a lower level) in adult SCs. *Myo6* mRNA is also detected in other non-HC populations (Kolla et al., 2020; Scheffer et al., 2015). Here, *Myo7a* was significantly enriched in adult OHCs (red arrows in Figure 6-figure supplement 1A) and also highly expressed in nascent HCs and OHC-like cells, but not in SCs that failed to become HCs (Figure 6B-C’’’). Therefore, we used *Myo7a* as an early HC marker to define the general HC fate in the RNA-Seq analysis described below.

### OHC-like cells globally upregulated wild-type adult OHC-enriched genes and downregulated adult SC-enriched genes

We next focused on characterizing OHC-like cells and determining the degree to which these were divergent from the original wild-type adult cochlear SCs. We manually picked 42 Tdtomato+ cells from *Fgfr3-Atoh1-Ikzf2-DTR* mice at P60 (Figure 6A) and sorted the cells into three types: (1) Tdtomato+/Myo7a+/Prestin+ cells (OHC-like cells, arrows in Figure 6C-C’’’; n = 11 cells); (2) Tdtomato+/Myo7a+/Prestin-cells (nascent HCs, asterisks in inset of Figure 6C-C’’’; n = 16); and (3) Tdtomato+/Myo7a-/Prestin-cells (defined as SCs that failed to become HCs, arrowheads in Figure 6C-C’’’; n = 15). The proportion of OHC-like cells was 26.2% (11/42), which agreed with the calculation (16.2%–29.0%) from immunostaining assays (Figure 4H). The result further validated the suitability of this criterion for our analysis. Moreover, expression of *Atoh1* and *Ikzf2* (arrows in Figure 6-figure supplement 2A) was enriched in all 42 cells, but not in wild-type adult SCs, which also confirmed that *Atoh1* and *Ikzf2* were permanently overexpressed in the 42 Tdtomato+ cells (regardless of their cell fates).

The results of both UMAP (uniform manifold approximation and projection) analysis and Pearson correlation-coefficient analysis demonstrated that, as compared with SCs that failed to become HCs, OHC-like cells were generally more divergent from adult wild-type SCs and convergent with adult wild-type OHCs (Figure 6D and E). Besides *Prestin* (*Slc26a5*) and *Myo7a* (arrows in Figure 6-figure supplement 2A), 1737 genes were expressed at a significantly higher level in OHC-like cells than in adult SCs. GO analysis of these 1737 genes revealed that the genes were enriched in functions involved in sound perception, cell differentiation, and inner ear development (Figure 6-figure supplement 2B), which further supported the notion that OHC-like cells globally behave like HCs. Notably, 824 out of the 1737 genes overlapped with genes enriched in wild-type adult OHCs, such as *Myo7a*, *Pvalb*, *Calb1*, *Rbm24*, *Cib2*, and *Lhfpl5* (arrows in Figure 6-figure supplement 2A). Conversely, 900 genes were expressed at significantly higher levels in adult wild-type SCs than in OHC-like cells, and 520 out of the 900 genes overlapped with adult wild-type SC-enriched genes, such as *Ceacam16*, *Bace2*, *Tuba1b*, *Gjb2*, and *Rorb* (arrows in Figure 6-figure supplement 2A). Supplemental File 4 contained the entire list of genes that were differently expressed in OHC-like cells and adult wild-type SCs, as well as the genes that overlapped with adult wild-type OHC- or SC-enriched genes.

In our examination of the difference between OHC-like cells and adult SCs, we also included nascent HCs and SCs that failed to become HCs as references for intermediate cell types. As expected, the identified genes were generally either not or only slightly unregulated/downregulated in SCs that failed to become HCs (Figure 6-figure supplement 2). Unexpectedly, however, we found that OHC-like cells and nascent HCs were similar to each other overall, and these were intermingled in the Pearson correlation-coefficient analysis (Figure 6E). This finding highlighted the heterogeneous gene-expression profiles among OHC-like cells and nascent HCs. Thus, although *Prestin* expression was detected in OHC-like cells but not in nascent HCs, the expression patterns of other HC genes might be the opposite. This possibility was partially supported by the finding that not all Prestin+ OHC-like cells expressed Rbm24, Pvalb, and Calb1, according to the results of both RNA-Seq and immunostaining assays (Figure 6-figure supplement 2 and Figure 6-figure supplement 3). *Rbm24*, *Pvalb*, and *Calb1* are early pan-HC markers and are normally turned on earlier than *Prestin* in wild-type OHCs (Grifone et al., 2018; Li et al., 2018; Liu et al., 2012a). Together, our results showed that OHC-like cells/nascent HCs, relative to SCs that failed to become HCs, were considerably more similar (but not identical) to adult wild-type OHCs. We next determined the developmental status of the wild-type OHCs to which the OHC-like cells were most similar.

### OHC-like cells were most similar to wild-type differentiating OHCs at neonatal ages

By reanalyzing raw data from a recent single-cell RNA-Seq study covering wild-type cochlear OHCs, IHCs, SCs, GER (greater epithelial ridge) cells, and LER (lesser epithelial ridge) cells (Kolla et al., 2020), we initially sorted out 87 OHCs at E16, 170 OHCs at P1, and 39 OHCs at P7 (Figure 6-figure supplement 4A). Because OHCs were highly heterogeneous due to the developmental basal-to-apical, medial-to-lateral gradient in the cochlear duct, trajectory analysis by using Monocle was applied to OHCs at the three aforementioned ages (Figure 6-figure supplement 4B and B’). Ultimately, we selected 59/87, 118/170, and 37/39 OHCs at E16, P1, and P7, respectively, and these cells either represented the majority or were located in the center of the cell populations at each age (within dotted lines in Figure 6-figure supplement 4 B). Lastly, the selected OHCs at the three ages were pooled with OHC-like cells, nascent HCs, P30_WT OHCs, and P60_WT SCs. Five main clusters were identified (Figure 6F): 10/11 OHC-like cells, 13/16 nascent HCs, and the majority of wild-type OHCs at P1 (P1_WT OHCs) belonged to cluster 1 (Figure 6G), whereas 1/11 OHC-like cells, 3/16 nascent HCs, and wild-type OHCs at E16 (E16_WT OHCs) formed cluster 2, which was close to cluster 5 (P60_WT SCs). Wild-type OHCs at P7 (P7_WT OHCs) and P30_WT OHCs formed cluster 3, which suggested that P7 OHCs were well differentiated (Jeng et al., 2020), and cluster 4 contained a small fraction of P1_WT OHCs. Thus, we concluded that OHC-like cells and nascent HCs were not distinguished at the transcriptomic level, and that these cells together were most similar to P1_WT OHCs. This conclusion was further supported by the presence of *Insm1* mRNA and protein in OHC-like cells (Figure 6-figure supplement 2A and Figure 6-figure supplement 3A-B’’’). Insm1 is only transiently expressed in differentiating OHCs at late embryonic or perinatal ages in a basal-to-apical gradient (Lorenzen et al., 2015).

### OHC-like cells were considerably less differentiated than adult wild-type OHCs

The finding that OHC-like cells were most similar to P1_WT OHCs led us to further compare the transcriptomic profiles between OHC-like cells and P30_WT OHCs and thus determine the main differences in their molecular signatures (Figure 6-figure supplement 5A). Here, the expression of *Atoh1*, but not *Ikzf2*, was higher in OHC-like cells than in P30_WT OHCs (Figure 6-figure supplement 5A); this was because *Atoh1* is not expressed in adult OHCs but Ikzf2 is (Chessum et al., 2018; Liu et al., 2014). Besides *Insm1*, *Hes6* was enriched in OHC-like cells (Figure 6-figure supplement 5A). Two previous reports have suggested that *Hes6* is expressed in cochlear HCs and is a target of Atoh1 (Qian et al., 2006; Scheffer et al., 2007). We also noted that adult wild-type SC-enriched genes such as *Fgfr3*, *Id2*, and *Id3* were expressed at higher levels in OHC-like cells than in P30_WT OHCs (Figure 6-figure supplement 5A); this was because these SC-enriched genes were not drastically downregulated in OHC-like cells. In Supplemental File 6, we summarized the entire list of genes differently expressed in OHC-like cells and P30_WT OHCs. GO analysis results showed that genes expressed at higher levels in OHC-like cells were enriched in cell adhesion, angiogenesis, and neuron projection development (Figure 6-figure supplement 5B). The enrichment of neural developmental genes in OHC-like cells was as expected: these genes are transiently expressed in wild-type differentiating HCs, but are gradually repressed by *Gfi1* in mature HCs (Matern et al., 2020). The complete gene list of each GO category was summarized in Supplemental File 7.

Conversely, genes such as *Ocm*, *Prestin (Slc26a5)*, *Lmo7*, and *Tmc1* were expressed at a lower level in OHC-like cells than in P30_WT OHCs (Figure 6-figure supplement 5A). These genes were enriched in adult wild-type OHCs (Supplemental File 1). Notably, *Lmo7* mutant mice have been reported to show abnormalities in HC stereocilia (Du et al., 2019). GO analysis revealed that the genes expressed at higher levels in P30_WT OHCs were enriched in transport and lipid transport (Figure 6-figure supplement 5C). The complete gene list of each GO category was summarized in Supplemental File 8.

## DISCUSSION

Our *in vivo* study clearly demonstrated the ability of *Ikzf2* and *Atoh1* to effectively convert adult cochlear SCs (mainly PCs and DCs) into Prestin+ OHC-like cells under the condition of OHC damage. Because adult cochlear SCs are considerably more challenging to reprogram than the corresponding neonatal cells, we considered our work to represent a notable advance in OHC regeneration studies. We expect *Atoh1* and *Ikzf2* to serve as potential targets for OHC regeneration in the clinic.

### Potential roles of Atoh1 and Ikzf2 in cell-fate transition from adult cochlear SCs into OHC-like cells

During normal development of HCs, Atoh1 is expressed for approximately one week, and the turning on/off of the expression follows a basal-to-apical gradient. Thus, the earlier Atoh1 expression is turned on at a specific location, the earlier it is turned off. During this period, Atoh1 is reported to perform at least three age-dependent functions: specifying the general HC fate, maintaining the survival of HC progenitors (short-term) and OHCs (long-term), and organizing the HC bundle (Bermingham et al., 1999; Cai et al., 2013; Woods et al., 2004). Although other functions cannot be ruled out, we speculated that the role of Atoh1 was to specify PCs and DCs with a general HC fate, which was partially evidenced by the inability of Ikzf2 alone to convert adult SCs into HCs. However, why Atoh1 alone was unable to trigger this conversion is unclear. The detailed synergistic effects between Atoh1 and Ikzf2 in the cell-fate conversion process warrants future investigation.

Ikzf2 is necessary for OHC maturation, and an Ikzf2 point-mutation model, *Ikzf2 ^cello/cello^*, displays early-onset hearing loss and diminished Prestin expression; by contrast, ectopic Ikzf2 induces Prestin expression in IHCs (Chessum et al., 2018). Whether *Prestin* is a direct target of Ikzf2 is unclear, but the onset of Ikzf2 expression at neonatal ages and the specific and permanent expression of the gene in OHCs suggest that Ikzf2 is a key TF required to specify and maintain the OHC fate, or to repress the IHC fate. Therefore, we speculated that the primary role of Ikzf2 was to direct nascent HCs into the OHC differentiation track, and, accordingly, Ikzf2 alone was able to induce Prestin expression in wild-type IHCs but not adult SCs, partially because IHCs are in the general HC differentiation track and are poised to turn on OHC genes. We also noted that the cell-fate switching from adult SCs to HCs induced by *Atoh1* and *Ikzf2* together occurred in a shorted period (∼12 days) than the switching from neonatal SCs to HCs induced by *Atoh1* alone (∼3 weeks) (Liu et al., 2012a). This again might result from the synergistic actions of Atoh1 and Ikzf2.

### Transcriptomic difference between Prestin+ OHC-like cells and adult wild-type OHCs

A promising advance reported here is our successful *in vivo* conversion of adult cochlear SCs into OHC-like cells expressing *Prestin*, *Insm1*, and *Ocm*, which are primarily expressed in OHCs but not IHCs (Lorenzen et al., 2015; Simmons et al., 2010; Zheng et al., 2000; Zhu et al., 2019); this reprogramming efficiency ranged between 16.2% and 29.0% depending on the different cochlear turns (Figure 4H). OHC-like cells upregulated 824 genes and downregulated 520 genes enriched in adult wild-type OHCs and SCs, respectively; however, the OHC-like cells differed from the fully differentiated OHCs present at P30 in both molecular and morphological aspects (Figures 5 and 6). For the analyses, we selected OHCs at P30 but not P60 because the OHC-like cells were derived from SCs at P30 and thus their intrinsic age might be ∼P30, although the mice were analyzed at P60. Notably, cochlear development is complete by P30, and wild-type OHCs at P30 and P60 are expected to differ minimally, a notion supported at least partly by the finding that adult cochlear SCs at P60 and P120 were indistinguishable from each other (Hoa et al., 2020).

To precisely evaluate the differentiation status of the cells studied here, we also reanalyzed data from a recent single-cell RNA-Seq study (Kolla et al., 2020). HC differentiation occurs in a basal-to-apical, medial-to-lateral gradient (Groves et al., 2013; Wu and Kelley, 2012), and to minimize gene-profiling variations among the differentiating OHCs at different cochlear turns, we selected OHCs at E16 that were less differentiated in the trajectory line, and these were assumed to be from the middle turn (Figure 6-figure supplement 4B). The non-selected OHCs at E16 were expected to be from the basal turn, and these were close to OHCs at P1. No apical OHCs were present at E16. Similarly, we selected OHCs at P1 that were distributed in the center of the trajectory line (Figure 6-figure supplement 4B), and thus we assumed that these cells were OHCs at P1 at the middle turn. The OHCs at P7 were highly homogenous and we speculated that these were apical (less differentiated) OHCs that could effectively tolerate the procedures used for preparing single-cell suspensions. Accordingly, with the same protocols being used, sequencing data obtained at younger ages were found to be of higher quality than the data from P7, and performing the analyses at ages after P7 was challenging (Kolla et al., 2020). Our data support the view that, instead of resembling P30_WT OHCs, the OHC-like cells were most similar to P1_WT OHCs (Figure 6F and G). The age P1 here might be a rough estimate because more precise gene profiling of OHCs between P1 and P7 is not yet available.

### Potential approach to promote OHC-like cells to resemble wild-type adult OHCs

What is the main disparity between the OHC-like cells described here and P30_WT OHCs? This is a key question because the final goal is to convert adult cochlear SCs into functional OHCs that exhibit high Prestin expression levels, overall molecular signatures, and cellular morphological features similar to those of wild-type adult OHCs. However, because hearing function was not rescued in this study (Figure 5D), additional methods must be used to further promote the current Insm1+ OHC-like cells expressing low levels of Prestin to move forward into the fully differentiated state in which the cells do not express Insm1 but abundantly express Prestin/Ocm (Lorenzen et al., 2015; Simmons et al., 2010; Zheng et al., 2000). Our results clearly showed that the stereocilium structure was one of the main differences between OHC-like cells and adult OHCs (Figure 5), but the OHC-like cells expressed multiple MET-channel-related proteins such as Cib2 and Tmc1 (Jia et al., 2020; Li et al., 2019; Pan et al., 2013). This agrees with the previous report that MET-channel assembly does not require normal hair-bundle morphology (Cai et al., 2013).

In future studies, we aim to focus on discovering additional critical genes such as *Emx2* that are necessary for hair-bundle organization and growth (Jacobo et al., 2019; Jiang et al., 2017). Instead of the OHC-like cells in our model here that were heterogeneous in terms of their gene profiles, we will compare neonatal OHCs at P1 in one particular cochlear duct location with their counterparts at P30; this should allow the prominent candidate genes that are differently expressed at the two ages to be sorted in a more efficient manner, and we will then employ our genetic loss-of-function screening approach (Zhang et al., 2018). Considering the promising results obtained such as finding the optimal phenotype of the OHC hair bundles being considerably shorter or more disorganized relative to control, but with the OHCs generated normally and surviving—we will combine additional candidate genes with *Atoh1* and *Ikzf2*, and this could lead to the generation of OHC-like cells that show superior differentiation relative to the cells produced here. In summary, we consider the *in vivo* ability of Atoh1 and Ikzf2 to successfully reprogram adult cochlear SCs into Prestin+ OHC-like cells in the presence of OHC damage to be a great advance and highly encouraging. We believe that Atoh1 and Ikzf2 will serve as key targets for future OHC regeneration therapies in the clinic.

## MATERIALS AND METHODS

### Generation of *Rosa26-CAG-LSL-Ikzf2/+* and *Prestin-DTR/+* knockin mouse strains by using CRISPR/Cas9 approach

The *Rosa26-CAG-Loxp-stop-Loxp-Ikzf2*3xHA-T2A-Tdtomato/+* (*Rosa26-CAG-LSL-Ikzf2/+*) knockin mouse strain was produced by co-injecting one sgRNA against the *Rosa26* locus (5ʹ-*ACTCCAGTCTTTCTAGAAGA*-3ʹ), donor DNA (Figure 1-figure supplement 1), and Cas9 mRNA into one-cell-stage mouse zygotes. A similar strategy was applied to generate the *Prestin-P2A-DTR/+* (*Prestin-DTR/+*) knockin mouse strain. The donor DNA is described in Figure 3-figure supplement 1, and the sgRNA against the *Prestin* (*Slc26a5*) locus was the following: 5ʹ-*CGAGGCATAAAGGCCCTGTA*-3ʹ. F0 mice with potential correct gene targeting were screened by performing junction PCR, and this was followed by crossing with wild-type C57BL/6 mice for germ-line transition (production of F1 mice). F1 mice were further confirmed by junction PCR. No random insertion of donor DNAs in the genome of F1 mice was detected in Southern blotting (Figure 1-figure supplement 1D and E, and Figure 3-figure supplement 1D and E), performed according to our previously described protocol (Li et al., 2018), and the two mouse strains were PCR-genotyped using tail DNA (representative gel images of PCR products were presented in Figure 1-figure supplement 1F and Figure 3-figure supplement 1F). Detailed primer sequences were described in Supplemental File 9. All mice were bred and raised in SPF-level animal rooms, and animal procedures were performed according to the guidelines (NA-032-2019) of the IACUC of Institute of Neuroscience (ION), CAS Center for Excellence in Brain Science and Intelligence Technology, Chinese Academy of Sciences.

### Sample processing, histology and immunofluorescence assays, and cell counting

Adult mice were anesthetized and sacrificed with the heart being perfused with 1× PBS and fresh 4% paraformaldehyde (PFA) to completely remove blood from the inner ear, after which inner ear tissues were dissected out carefully, post-fixed with fresh 4% PFA overnight at 4°C, and washed thrice with 1× PBS. The adult inner ears were first were decalcified with 120 mM EDTA for 2 days at 4°C until they were soft and ready for micro-dissecting out the cochlear sensory epithelium for use in whole-amount preparation and immunostaining with the following first antibodies: anti-HA (rat, 1:200, 11867423001, Roche), anti-Prestin (goat, 1:1000, sc-22692, Santa Cruz), anti-Myo6 (rabbit, 1:500, 25-6791, Proteus Bioscience), anti-Myo7a (rabbit, 1:500, 25-6791, Proteus Bioscience), anti-Sox2 (goat, 1:500, sc-17320, Santa Cruz), anti-Insm1 (guinea pig, 1:6000; a kind gift from Dr. Shiqi Jia from Jinan University, Guangzhou, China, and Dr. Carmen Birchmeier from Max Delbrück Center for Molecular Medicine, Berlin, Germany), anti-Parvalbumin (mouse, 1:500, P3088, Sigma), anti-Rbm24 (rabbit, 1:500, 18178-1-AP, Proteintech), anti-Calbindin (rabbit, 1:500, C9848, Sigma), and anti-vGlut3 (rabbit, 1:500, 135203, Synaptic System). Cochlear tissues were counterstained with Hoechst 33342 solution in PBST (1:1000, 62249, Thermo Scientific) to visualize nuclei, and were mounted with Prolong gold antifade medium (P36930, Thermo Scientific). Nikon C2, TiE-A1, and NiE-A1 plus confocal microscopes were used to capture images.

Each whole-mount preparation of the cochlear duct was divided into three parts, which were initially scanned at 10× magnification under a confocal microscope. In the obtained images, a line was drawn in the middle of IHCs and OHCs to precisely measure the entire length of each cochlear duct, which was then divided into basal, middle, and apical portions that were of equal length. For determining the degree of OHC death in the *Prestin-DTR/+* model (Figure 3), in each mouse, both the OHC and the IHC numbers in the same scanning regions (60×, confocal microscopy) were quantified at each turn (two different areas were selected and the average number was calculated); the IHC number was used as a reference because IHCs did not die following DT treatment. In terms of counting Prestin+ IHCs in *Atoh1-CreER+; Rosa26-CAG-LSL-Ikzf2/+* mice (Figure 1D), and Tdtomato+/HA+ cells, OHC-like cells, and nascent HCs in *Fgfr3-Atoh1-Ikzf2* and *Fgfr3-Atoh1-Ikzf2-DTR* mice (Figs. 2 and 4), the entire cochlear duct of each mouse was scanned using a confocal microscope (60×) to minimize variation between different replicates. All cell counting data are presented as means ± SEM. Statistical analyses were performed using one-way ANOVA, followed by a Student’s *t* test with Bonferroni correction. GraphPad Prism 6.0 was used for all statistical analyses.

### Single-cell RNA-Seq and bioinformatics analysis

Three different models were used to pick Tdtomato+ cells: (1) *Prestin-CreER/+; Ai9/+* mice that were injected with tamoxifen at P20 and P21; all Tdtomato+ cells were OHCs at P30, because of exclusive Cre activity in OHCs (Fang et al., 2012). (2) *Fgfr3-iCreER+; Ai9/+* mice that were administered tamoxifen at P30 and P31; all Tdtomato+ cells within the cochlear sensory epithelium were SCs (primarily PCs and DCs) at P60, according to our previous reports (Liu et al., 2012a; Liu et al., 2012b). (3) *Fgfr3-Atoh1-Ikzf2-DTR* mice that were administered tamoxifen at P30 and P31 and then DT at P36. The Tdtomato+ cells within the cochlear sensory epithelium at P60 included OHC-like cells, nascent HCs, and SCs that failed to become HCs.

The cochlear sensory epithelium from each model mouse was carefully dissected out, digested, and used for preparing single-cell suspensions according to our detailed protocol described previously (Li et al., 2020a). All the aforementioned Tdtomato+ cells were picked under a fluorescence microscope (M205FA, Leica) as described in Figure 6A. We picked 17 wild-type adult OHCs at P30, 16 wild-type adult SCs at P60, and 42 Tdtomato+ cells from the *Fgfr3-Atoh1-Ikzf2-DTR* model (Figure 6D), and we immediately used these cells for reverse-transcription and cDNA amplification with a Smart-Seq HT kit (Cat# 634437, Takara). The cDNAs (1 ng each) were tagmented using a TruePrep DNA Library Prep Kit V2 for Illumina (Cat# TD503, Vazyme) and a TruePrep Index Kit V2 for Illumina (Cat# TD202, Vazyme). The final libraries were subject to paired-end sequencing on the Illumina Novaseq platform. Each library was sequenced to obtain 4G raw data.

FastQC (v0.11.9) and trimmomatic (v0.39) were used for quality control of raw sequencing data. For ∼70% -80% of the reads, high-quality mapping to the mouse reference genome (GRCm38) was achieved by using Hisat2 (v2.1.0) with default parameters. Raw counts were calculated using HTSeq (v0.10.0), and gene-expression levels were estimated by using StringTie (v1.3.5) with default parameters. Gene abundances were presented as transcript per million (TPM) values. Differentially expressed genes (DEGs) were analyzed using R package “DESeq2” (p.adj < 0.05, absolute value of (log2 Fold Change) > 2), and the DEGs were used for determining biological process enrichment (p < 0.05, adjusted using Bonferroni correction) by using DAVID (Database for Annotation, Visualization and Integrated Discovery). All the raw data from our single-cell RNA-Seq work have been deposited in the GEO (Gene Expression Omnibus) under Accession No. GSE161156.

Seurat (R package v3.0) was applied to the raw data collected for wild-type OHCs at E16, P1, and P7 in a recent study (Kolla et al., 2020). To more precisely compare the transcriptomic profiles obtained from 10× genomics and smartseq approaches, we integrated them first by using the functions “FindIntegrationAnchors” (k.filter = 30) and “IntegrateData.” Principal components (PCs) were calculated using the “RunPCA” function, and the top 20 PCs were used for the dimensionality reduction process (“RunTSNE” and “RunUMAP”). Unsupervised clustering was performed using the “FindClusters” function (resolution = 0.5). Furthermore, to more precisely define OHCs (E16, E17, and P7) versus OHC progenitors (E14), we used criteria that were more stringent than those applied in the previous report. E16_WT OHCs were defined as cells in which the individual expression level of *Insm1*, *Myo6*, and *Atoh1* was above zero; in P1_WT OHCs, *Bcl11b*, *Myo6*, *Myo7a*, and *Atoh1* expression was above zero; and in P7_WT OHCs, *Prestin* (*Slc26a5*), *Myo6*, *Ocm*, and *Ikzf2* expression was above zero. Trajectory analysis was performed using Monocle (R package v2.0). Pre-processed Seurat datasets were imported into Monocle by using the “importCDS” function. The results of differential expression (DE) analysis identified genes that were significantly altered among OHCs at the three ages. Cells were ordered along a pseudotime axis by using the “orderCells” function in Monocle.

### ABR measurement

ABR measurements were performed by using sound at 4k, 5.6k, 8k, 11.3k, 16k, 22.6k, and 32k Hz, according to the detailed protocol described in our previous report (Li et al., 2018). Student’s *t* test was used for statistical analysis at each frequency between two conditions (Figure 3C and Figure 5D).

### Tamoxifen and DT treatment

Tamoxifen (Cat# T5648, Sigma) was dissolved in corn oil (Cat# C8267, Sigma) and injected intraperitoneally at 3 mg/40 g body weight (P0 and P1) or 9 mg/40 g body weight (P20 and P21, and P30 and P31). DT (Cat# D0564, Sigma) dissolved in 0.9% NaCl solution was also delivered through intraperitoneal injection, at a dose of 20 ng/g body weight, and the mice were analyzed at P42 or P46 or P60.

### SEM analysis

SEM was performed following the protocol reported previously (Parker et al., 2016). Briefly, we made holes at the cochlear apex and then washed the samples gently with 0.9% NaCl (Cat#10019318, Sinopharm Chemical Reagent Co, Ltd.) and fixed them with 2.5% glutaraldehyde (Cat# G5882, Sigma) overnight at 4°C. On the following day, the cochlear samples were washed thrice with 1× PBS and then decalcified using 10% EDTA (Cat# ST066, Beyotime) for 1 day, post-fixed for 1 h with 1% osmium tetroxide (Cat#18451, Ted Pella), and subject to a second fixation for 30 min with thiocarbohydrazide (Cat#88535, Sigma) and a third fixation for 1 h with osmium tetroxide. Next, the cochlear samples were dehydrated using a graded ethanol series (30%, 50%, 75%, 80%, 95%; Cat#10009259, Sinopharm Chemical Reagent Co, Ltd.), at 4°C with 30 min used at each step, and, lastly, dehydrated in 100% ethanol thrice (30 min each) at 4°C. The cochlear samples were dried in a critical point dryer (Model: EM CPD300, Leica), after which whole-mount cochlear samples were prepared under a microscope to ensure that hair bundles were facing upward and then treated in a turbomolecular pumped coater (Model: Q150T ES, Quorum). The final prepared samples were scanned using a field-emission SEM instrument (Model: GeminiSEM 300, ZEISS).

## ACKNOWLEDGMENTS

We thank Drs. Qian Hu, Yu Kong, and Xu Wang from Optical Imaging and EM Facility of the Institute of Neuroscience (ION) for support with image analysis; Dr. Hui Yang (Principal Investigator at the ION) for sharing the zygote microinjection system used to generate the knockin mice; Ms. Qian Liu (from the Department of Embryology of the ION animal center) for helping us in transplanting zygotes into pseudopregnant female mice; and Dr. Shiqi Jia (Jinan University, Guangzhou, China) and Dr. Carmen Birchmeier (Max Delbrück Center for Molecular Medicine, Berlin, Germany) for kindly providing the anti-Insm1 antibody.

## SUPPLEMENTAL FIGURE LEGENDS

**Figure 1-figure supplement 1.**
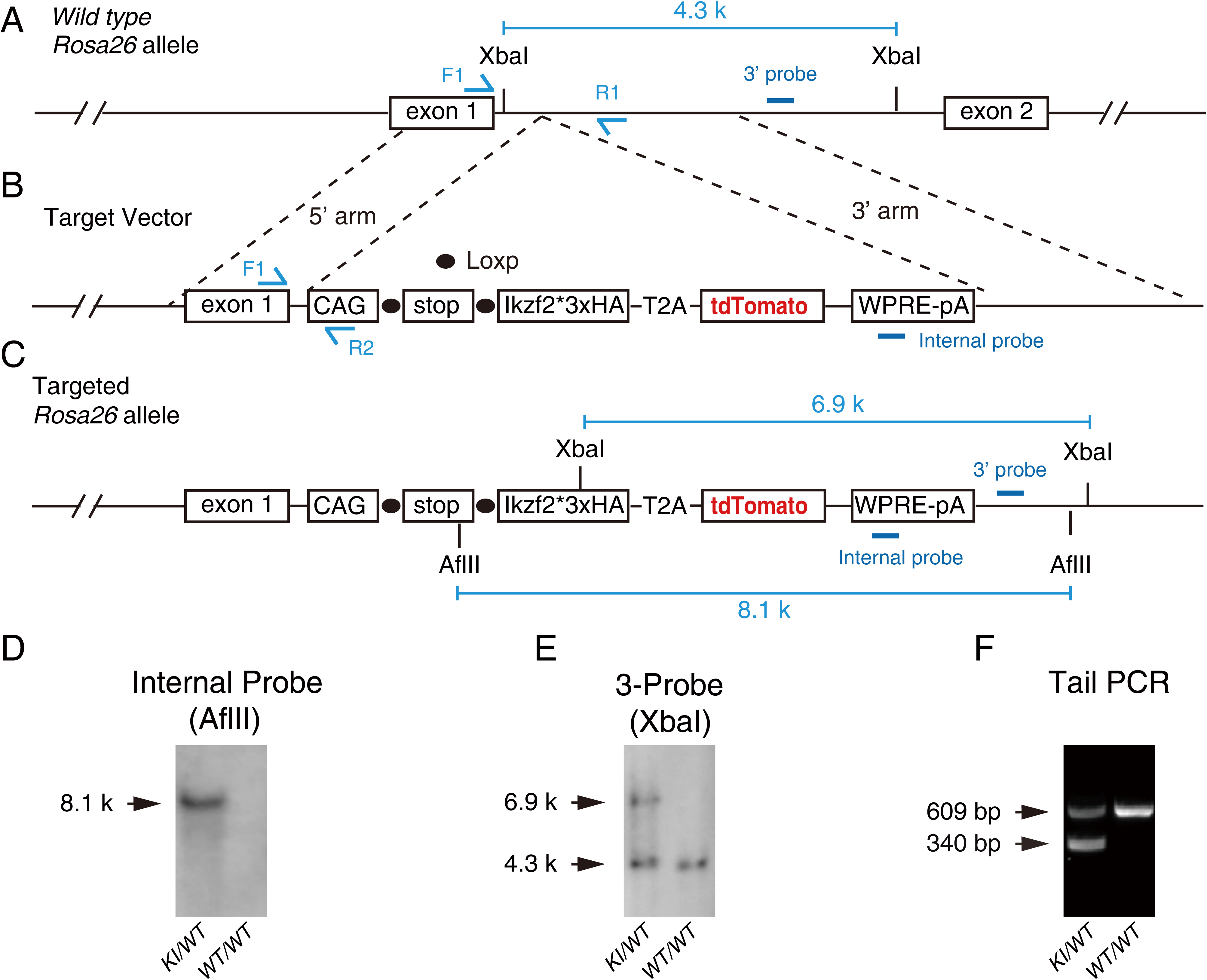
Generation of *Rosa26-CAG-LSL-Ikzf2/+* knockin mouse model by using CRISPR/Cas9 method. **(A)** Wild-type *Rosa26* allele. **(B)** Illustration of target vector. Ikzf2 was tagged with 3× HA fragments at its C-terminus, and this was followed by T2A-Tdtomato; Ikzf2 and Tdtomato were transcribed and translated together as a fusion protein, but then cleaved into Ikzf2 and Tdtomato, respectively, through the 2A approach. **(C)** Illustration of *Rosa26* after correct gene targeting. **(D-E)** Southern blotting assay of internal probe (D) and 3ʹ-end probe (E). **(F)** Genotyping PCR of tail DNA extracted from heterozygous (KI/WT) and wild-type (WT/WT) mice. WT mice showed a single 609-bp band, whereas heterozygous mice showed 609- and 340-bp bands.

**Figure 1-figure supplement 2.**
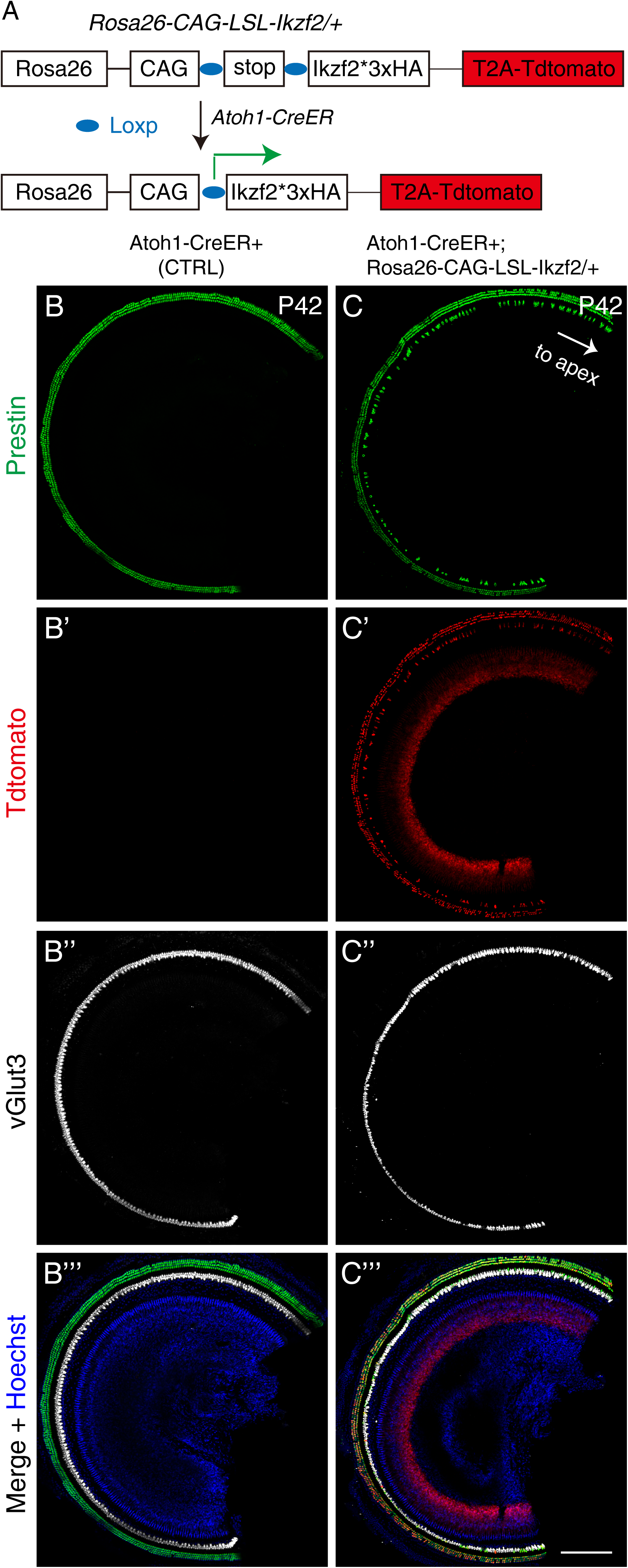
Prestin was expressed in IHCs with ectopic Ikzf2. **(A)** Same as illustration in Figure 1A. **(B-C’’’)** Triple labeling for Prestin, Tdtomato, and vGlut3 in both control *Atoh1-CreER+* mice (B-B’’’) and experimental *Atoh1-CreER+; Rosa26-CAG-LSL-Ikzf2/+* mice (C-C’’’) at P42; these low-magnification images compliment the images in Figure 1. Prestin was expressed in vGlut3+ IHCs of experimental but not control mice, and Prestin expression again completely overlapped with that of Tdtomato. Scale bar: 200 μm.

**Figure 3-figure supplement 1.**
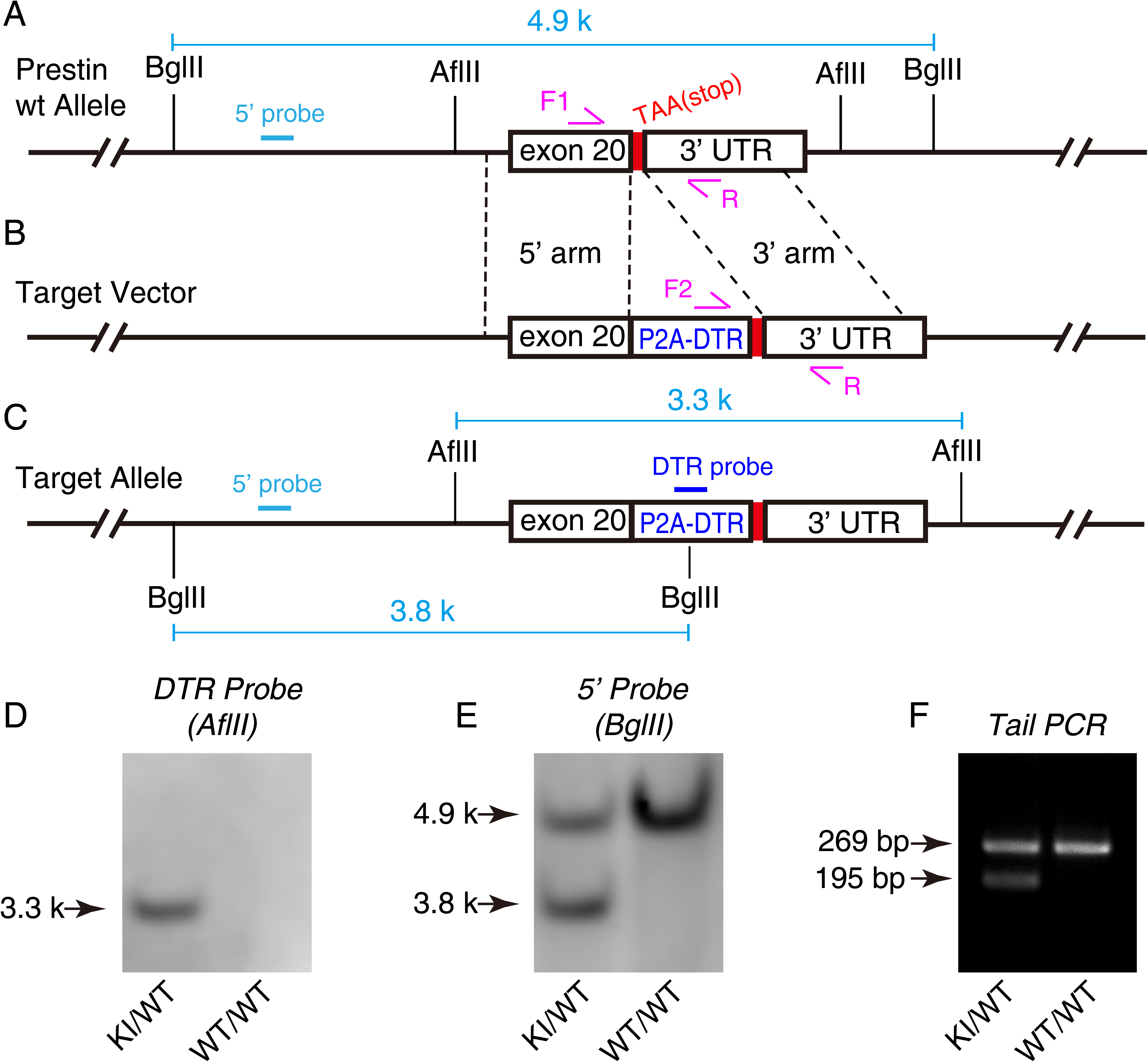
Generation of *Prestin-DTR/+* knockin mouse model. **(A)** Wild-type *Prestin* (*Slc26a5*) allele. **(B)** Illustration of target vector. P2A-DTR (diphtheria toxin receptor) was inserted immediately before the stop codon *TAA*. DTR expression was completely controlled by *Prestin* promoter and enhancer. **(C)** Illustration of *Prestin* allele after correct gene targeting. **(D-E)** Southern blotting assay of internal DTR probe (D) and 5ʹ-end probe (E). **(F)** Genotyping PCR of tail DNA extracted from heterozygous (KI/WT) and wild-type (WT/WT) mice. WT mice displayed a single 269-bp band, whereas heterozygous mice showed 195- and 269-bp bands.

**Figure 4-figure supplement 1.**
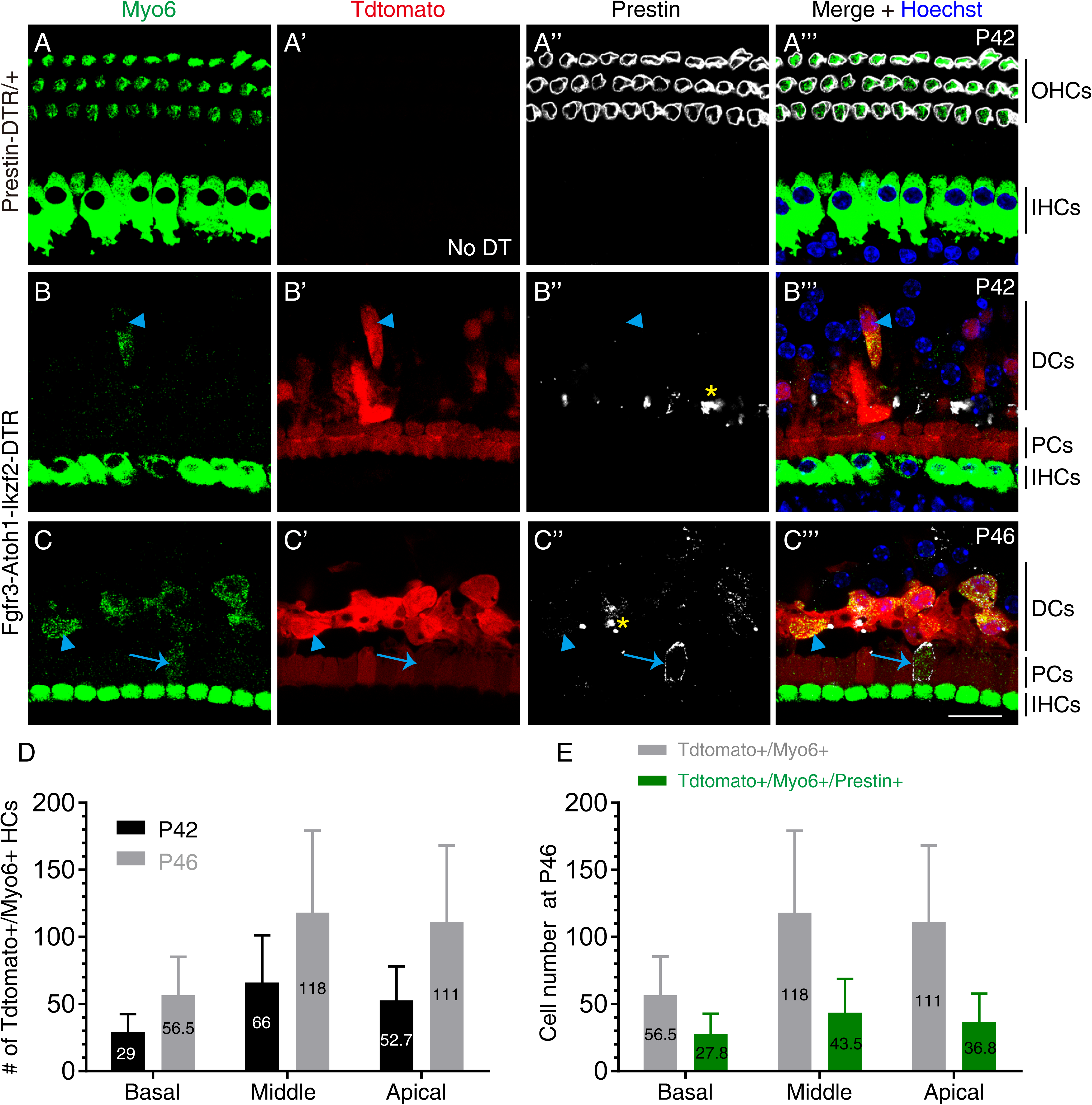
Nascent HCs emerged at P42 and OHC-like cells at P46. (A-C’’’) Triple labeling for early HC marker Myo6, Tdtomato, and Prestin in control mice at P42 (A-A’’’) and in experimental mice at P42 (B-B’’’) and P46 (C-C’’’). Arrowheads: nascent HCs that were Tdtomato+/Myo6+/Prestin-; arrows: OHC-like cells that were Tdtomato+/Myo6+/Prestin+; asterisks: debris of dying endogenous OHCs. **(D)** Quantification of new HCs at different cochlear turns in *Fgfr3-Atoh1-Ikzf2-DTR* mice at P42 and P46. More new HCs tended to be present at P46 than P42, although no statistical difference was calculated due to large variations in the numbers; mean numbers of nascent HCs are shown. **(E)** Quantification of new HCs (nascent and OHC-like cells together) and OHC-like cells at the same turn in *Fgfr3-Atoh1-Ikzf2-DTR* mice at P46; mean numbers of new HCs and OHC-like cells were shown. Scale bar: 20 μm.

**Figure 6-figure supplement 1.**
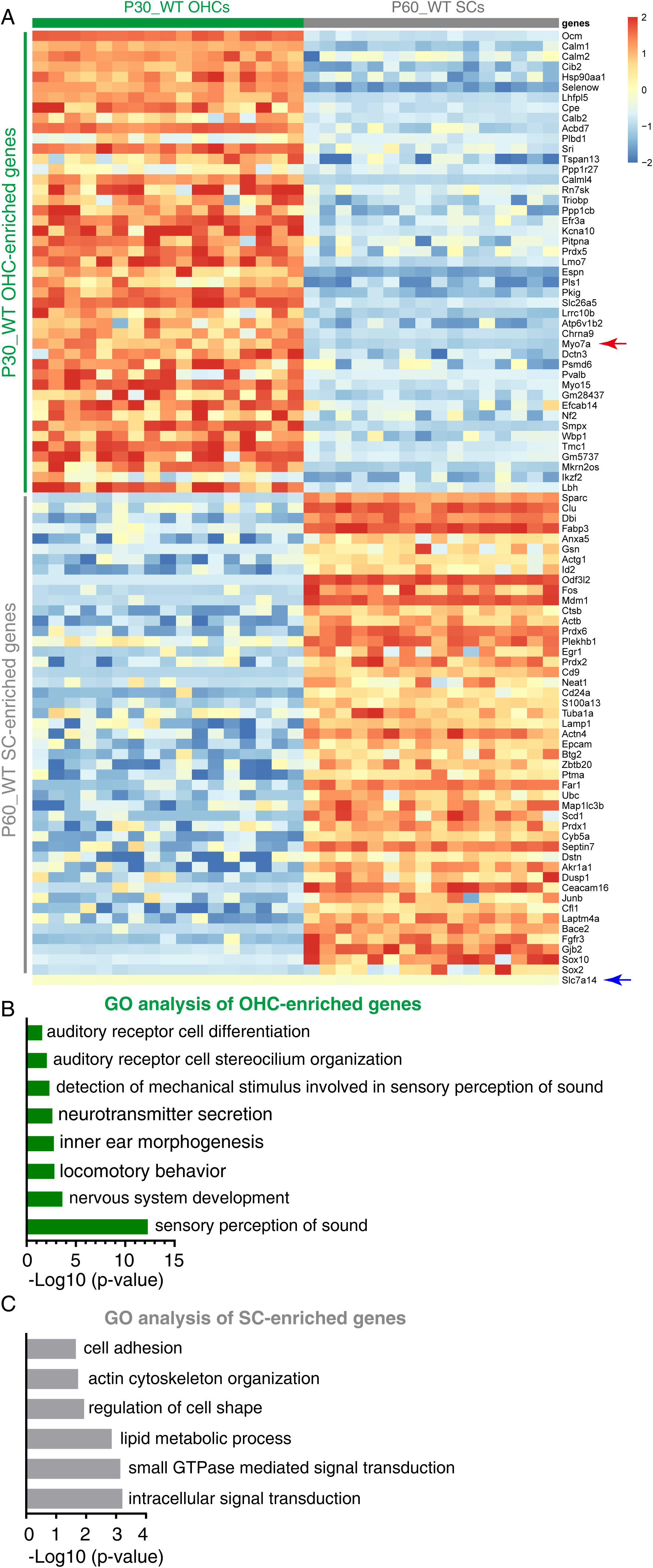
Transcriptomic comparison between adult wild-type OHCs and SCs. We manually picked 17 wild-type OHCs at P30 (P30_WT OHCs) and 16 SCs at P60 (P60_WT SCs) and performed single-cell RNA-Seq by using the smartseq approach. **(A)** Examples of top enriched genes in P30_WT OHCs and P60_WT SCs; the complete gene list appears in Supplemental File 1. Early pan-HC-specific genes such as *Myo7a* (red arrow) and OHC-specific genes such as *Ocm*, *Lbh*, *Prestin* (*Slc26a5*), and *Ikzf2* were incorporated with the genes enriched in P30_WT OHCs. Similarly, SC markers such as *Sox2*, *Sox10*, *Bace2*, and *Ceacam16* were included among the genes enriched in P60_WT SCs. As expected, the IHC-specific gene *Slc7a14* was included with neither OHC-nor SC-enriched genes. **(B-C)** Gene Ontology (GO) enrichment analysis of genes enriched in OHCs (B, green) and SCs (C, gray). The gene lists of each GO category appeared in Supplemental Files 2 and 3.

**Figure 6-figure supplement 2.**
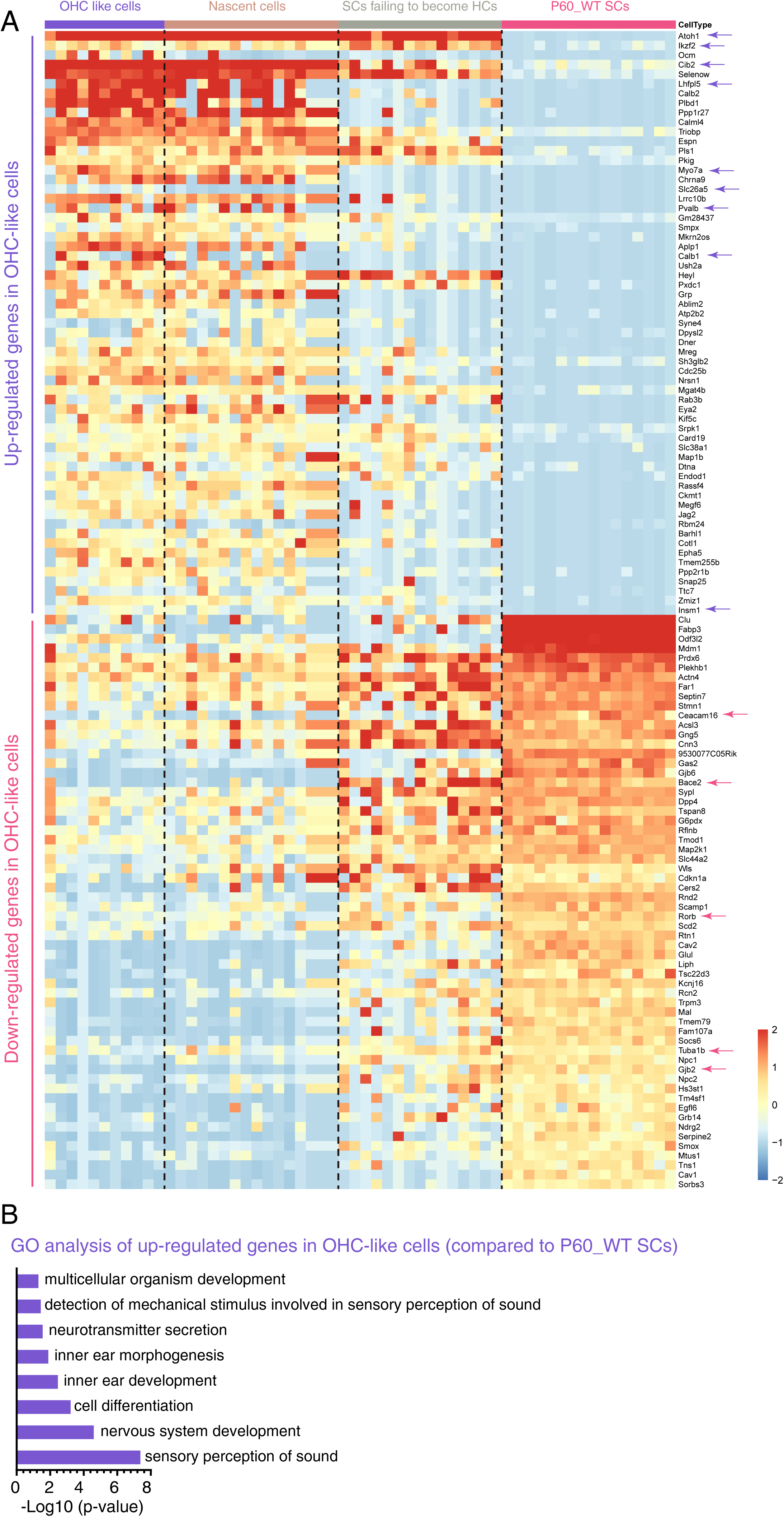
Hundreds OHC and SC genes were upregulated and downregulated, respectively, in OHC-like cells. Single-cell RNA-Seq was applied to 42 Tdtomato+ cells from *Fgfr3-Atoh1-Ikzf2-DTR* mice at P60; the cells were categorized as OHC-like cells, nascent HCs, and SCs that failed to become HCs. We compared the transcriptomic profiles between OHC-like cells and P60_WT SCs. **(A**) Relative to P60_WT SCs, top significantly upregulated or downregulated genes in OHC-like cells were sorted first and only those that overlapped with genes enriched in P30_WT OHCs or P60_WT SCs were selected and presented as examples. Here, nascent HCs and SCs that failed to become HCs were included as references only. The complete gene lists were presented in Supplemental File 4. **(B)** GO analysis of all genes (without using overlap with genes enriched in P30_WT OHCs) that were significantly upregulated in OHC-like cells. The gene lists of each GO category appeared in Supplemental File 5.

**Figure 6-figure supplement 3.**
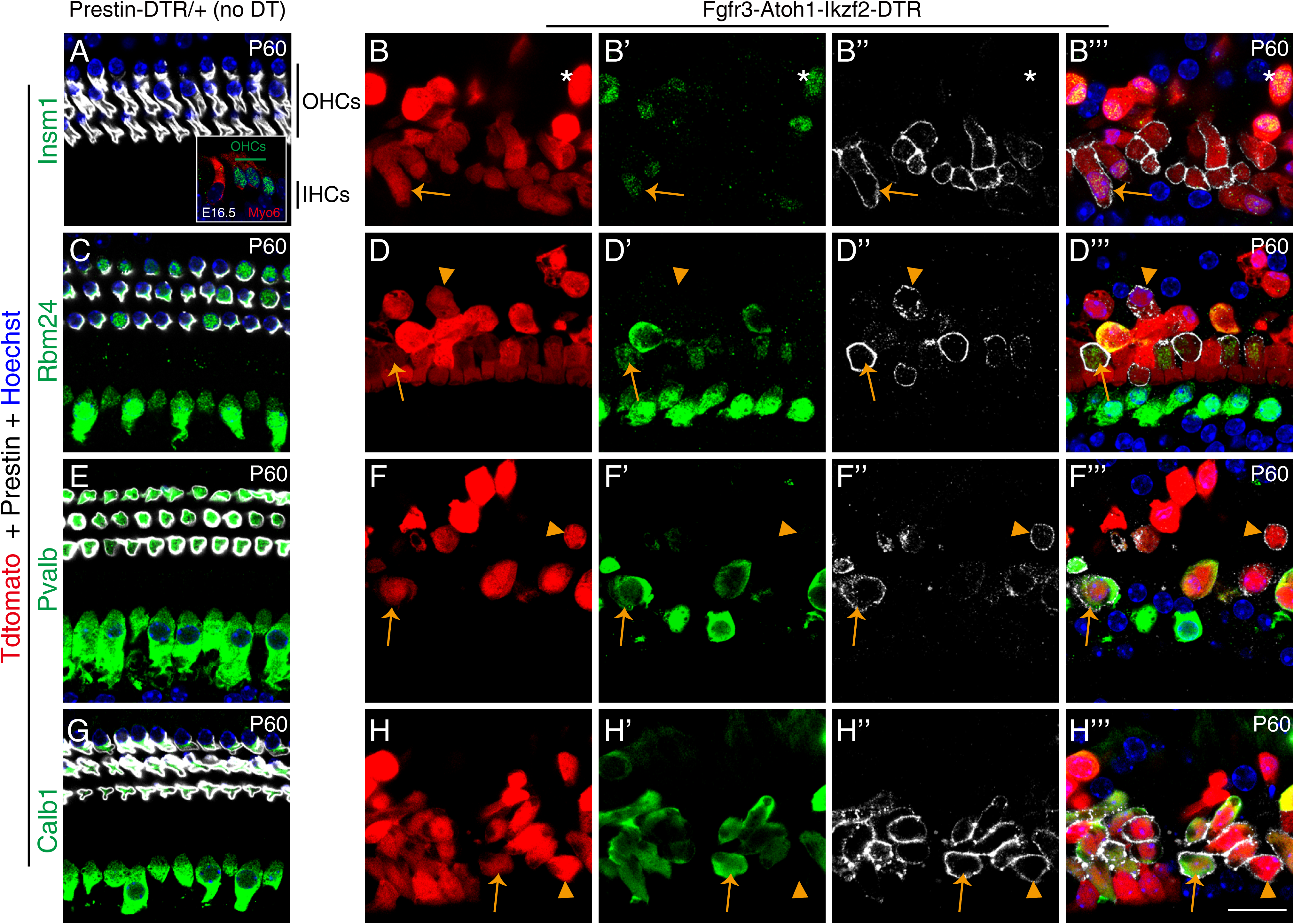
Heterogeneous expression of OHC genes in OHC-like cells. Cochlear samples (P60) from control *Prestin-DTR/+* mice not treated with DT (A, C, E, G) and experimental *Fgfr3-Atoh1-Ikzf2-DTR* mice treated with DT (B-B’’’, D-D’’’, F-F’’’, H-H’’’) were triple labeled for Tdtomato (red), Prestin (white in images), and a third different marker (antibody staining, shown in green): Insm1 (A-B’’’), Rbm24 (C-D’’’), Pvalb (E-F’’’), or Calb1 (G-H’’’). Asterisks in (B-B’’’): cell that was Tdtomato+/Insm1+ but did not yet express Prestin; arrows in (B-B’’’): cell that was Tdtomato+/Insm1+/Prestin+. Inset in (A): cochlear cryosection sample at E16.5 stained for Myo6 and Insm1. Insm1 expression was detected in OHCs but not in IHCs, which was presented as another control to confirm the specificity of the Insm1 antibody. Arrows in (D-D’’’, F-F’’’, H-H’’’): triple-positive cells; arrowheads in (D-D’’’, F-F’’’, H-H’’’): cells that were Tdtomato+/Prestin+ but did not express Rbm24 (D-D’’’), Pvalb (F-F’’’), or Calb1 (H-H’’’). Prestin in (B’’, D’’, F’’, H’’) was shown at a higher gain than in (A, C, E, G) to enhance signal visualization; the Prestin level in OHC-like cells was considerably lower than that in wild-type OHCs. Scale bar: 20 μm.

**Figure 6-figure supplement 4.**
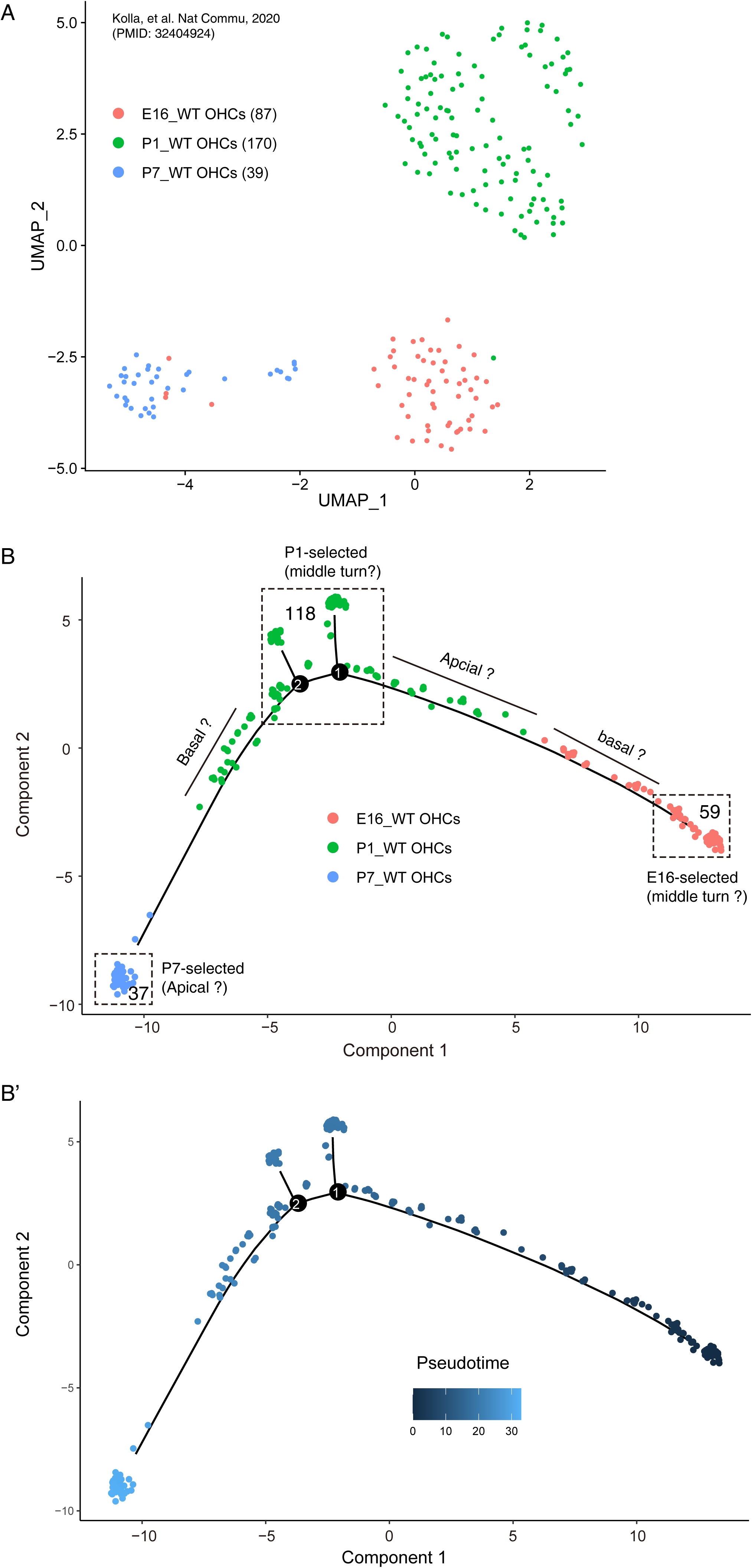
Trajectory analysis of wild type differentiating OHCs. **(A)** UMAP analysis of wild-type OHCs at E16, P1, and P7. OHCs of the same age formed their own main cluster. **(B-B’)** Trajectory analysis of OHCs at three different ages by using Monocle. Cells within dotted lines were selected for further analysis as described in Figure 6.

**Figure 6-figure supplement 5.**
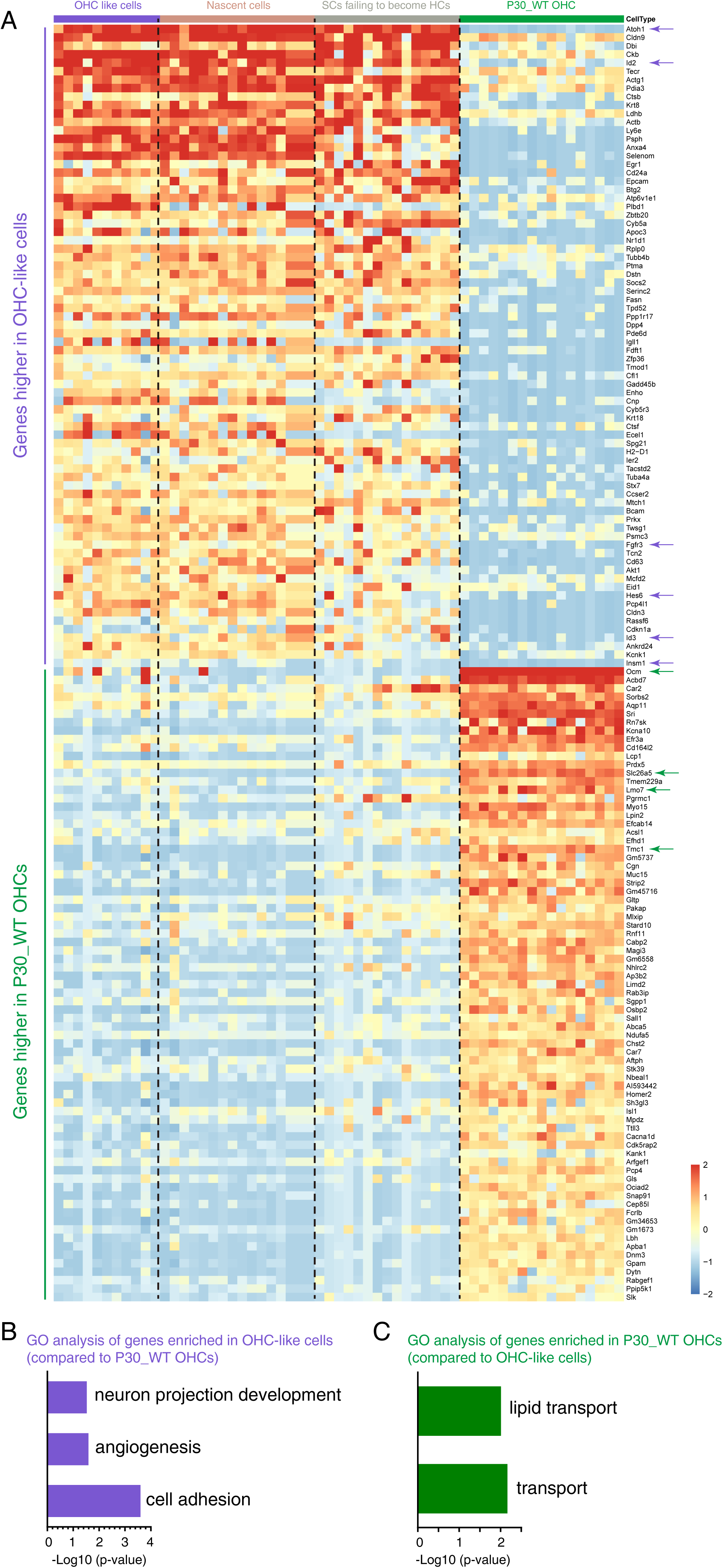
Transcriptomic difference between OHC-like cells and mature OHCs. **(A)** Examples of top DEGs between OHC-like cells and wild-type OHCs at P30 (P30_WT OHCs). Nascent HCs and SCs that failed to become HCs were included as references only. *Atoh1*, *Insm1*, and *Hes6*, which are transiently expressed in OHCs, were expressed at a higher level in OHC-like cells than in P30_WT OHCs. *Id2, Id3*, and *Fgfr3* are normally expressed in SCs but were not significantly decreased in OHC-like cells; these genes were indicated by purple arrows. Conversely, *Ocm*, *Prestin* (*Slc26a5*), *Lmo7*, and *Tmc1* showed increased expression in P30_WT OHCs, and these genes encode functional proteins in mature OHCs (green arrows). The complete gene lists were presented in Supplemental File 6. **(B-C)** GO analysis of genes upregulated in OHC-like cells (B) or P30_WT OHCs (C). The gene lists of each GO category were included in Supplemental Files 7 and 8, respectively.

